# Dopamine Builds and Reveals Reward-Associated Latent Behavioral Attractors

**DOI:** 10.1101/2022.07.07.499108

**Authors:** J. Naudé, MXB. Sarazin, S. Mondoloni, B. Hanesse, E. Vicq, F. Amegandjin, A. Mourot, P. Faure, B. Delord

**Affiliations:** Sorbonne Université, Inserm, CNRS, Neuroscience Paris Seine - Institut de biologie Paris Seine (NPS - IBPS), 75005 Paris, France; Institut des Systèmes Intelligents et de Robotique (ISIR), Sorbonne Université, CNRS, Paris, France; Brain Plasticity Unit, CNRS, ESPCI Paris, PSL Research University, 75005 Paris, France; CNRS, Université de Montpellier, INSERM - Institut de Génomique Fonctionnelle, 34094 Montpellier, France

## Abstract

Phasic variations in dopamine levels are interpreted as a teaching signal reinforcing rewarded behaviors. However, behavior also depends on the online, neuromodulatory effect of phasic dopamine signaling. Here, we unravel a new neurodynamical principle that reconciles these roles. In a biophysical recurrent network-based decision architecture, we showed that dopamine-mediated synaptic plasticity stabilized neural assemblies representing rewarded locations as latent, local attractors. Dopamine-modulated synaptic excitability activated these attractors online, and they became accessible as internal goals, even from remote animal positions. We experimentally validated these predictions in mice, using optogenetics, by demonstrating that online dopamine signaling specifically attracts animals toward rewarded locations, without off-target motor effects. We therefore propose that online dopamine signaling reveals potential goals by widening and deepening the basin of dopamine-built attractors representing rewards.

## INTRODUCTION

Transient, phasic dopamine (DA) release contributes both to learning (updating the values used to make future decisions based on experience) and to motivation (making ongoing decisions and invigorating goal-oriented behaviors), but reconciling these two roles within a unified theory of DA function has remained challenging (*1–3*). The popular reinforcement learning (RL) theory interprets phasic DA signaling uniquely as a reward-related teaching signal (*1, 4*), which functions by modulating long-term synaptic plasticity (*5*–*7*) to build neural representations of the value of actions that have previously led to reward (*8*, *9*). This role of DA in value learning is well demonstrated by the robust conditioned place preference induced by optogenetic stimulation of DA cells in the ventral tegmental area (VTA) (*10, 11*).

However, RL theory does not define, nor account for, a role for phasic DA signaling in ongoing behavior (*1*, *4*) despite renewed interest in the evidence linking this activity with motivation (*3*, *12*, *13*). Phasic DA neuron activity indeed occurs during self-paced movement initiation (*14–17*), and phasic optogenetic stimulation of DA neurons drives action initiation (*11*, *17 but see 16*). Accounts of these immediate effects of DA suggest either a “directional” role with DA signals specifying the decision to be taken (*18*) or an “activational” or energizing role, with DA determining the level of motor resources to engage in performing an action (*3*, *13, 19*). The limited encoding capacity of DA cells (*20*) and the larger impact of DA antagonist administration on action probability and vigor rather than on preferences, argues for an activational role of DA signaling (*19*). However, within this activational framework, DA would gate decision-making by lowering a universal decision threshold, increasing the probability and reducing the latency of all actions. It nevertheless remains unclear within these decision-threshold models how exactly DA induces movement energization. Furthermore, DA clearly does not have the same impact on all actions, which goes against decision-threshold models. DA signaling is mostly associated with, and necessary for, non-stereotyped, anticipatory, distal, or effortful behaviors, i.e. when some physical or cognitive distance separate the animal from a reward (*2*, *21*, *22*). Such a role of DA, which cannot be considered as purely activational nor as purely directional, is therefore still poorly explained by reinforcement learning theory.

Rather than deriving DA’s role from a phenomenological model of decision-making, we used dynamical systems theory to assess the biophysical effects of learning and motivational DA effects within a distributed decision architecture, which we called the MAGNet (Motivational Attraction toward Goals through NETwork dynamics) model. DA modulation of synaptic plasticity is believed to carve “Hebbian” assemblies (*23*) of strongly interconnected neurons, representing a decision that was repeatedly rewarded. Such Hebbian assemblies can be considered as attractors of network dynamics (*24*), i.e. particular states of sustained, reverberating activity (*25–28*) toward which the network activity converges. In standard models, convergence from a rest state toward the decision-related attractor either requires a cue stimulus (*26*), or is driven by noise (*27*). Here, we propose that the motivational role of phasic DA signaling is to reveal latent (i.e. not necessarily expressed) attractors previously built by DA-modulated plasticity, and to promote transitions from a resting state to engagement in decision-related attractor dynamics. By testing the MAGNet model with experimental data in a learned task, we demonstrate that, rather than increasing the probability of every action (*3*, *17*), phasic DA activity specifically gates previously learned goals. We thus reinterpret the motivational role of phasic DA signaling as a dynamic process that increases the accessibility of attractors representing potential goals.

## RESULTS

To characterize the role of phasic, transient dopamine (DA) signaling from the ventral tegmental area (VTA) in both reinforcement learning and motivation, we used an un-cued optogenetic conditioning task. Indeed, cues previously paired with reward induce both the release of DA and an increase in motor responses (*29*), confounding the interpretation of the role of DA in subsequent behavior. Instead, we designed a task which requires mice to learn an internal memory of a rewarded location. We achieved selective manipulation of dopamine neurons by expressing ChR2 in the VTA from dopamine transporter (DAT)-Cre mice (Figure 1a). We then placed mice into a circular open-field where we paired three explicit locations with 500 ms, 20 Hz VTA photostimulation (Figure 1a), which drives bursting activity in dopamine neurons (Supplementary Figure 1). As mice explored the open field, DA neurons were stimulated when mice were detected on one of the locations. Two consecutive visits to the same location were not paired with photostimulation, prompting mice to constantly alternate between the rewarded locations (*30*, *31*). Mice increased the number of photostimulations earned with learning sessions (Figure 1b, two-way ANOVA with repeated measures, groups: F_(1)_=30.04, p=0, time: F_(9)_=5.69, p=0, interaction: F_(1,9)_=3.9, p=0.0002), which confirmed that phasic bursting in VTA DA neurons constitutes a teaching signal for place–reward association (*32*).

**Figure 1.**
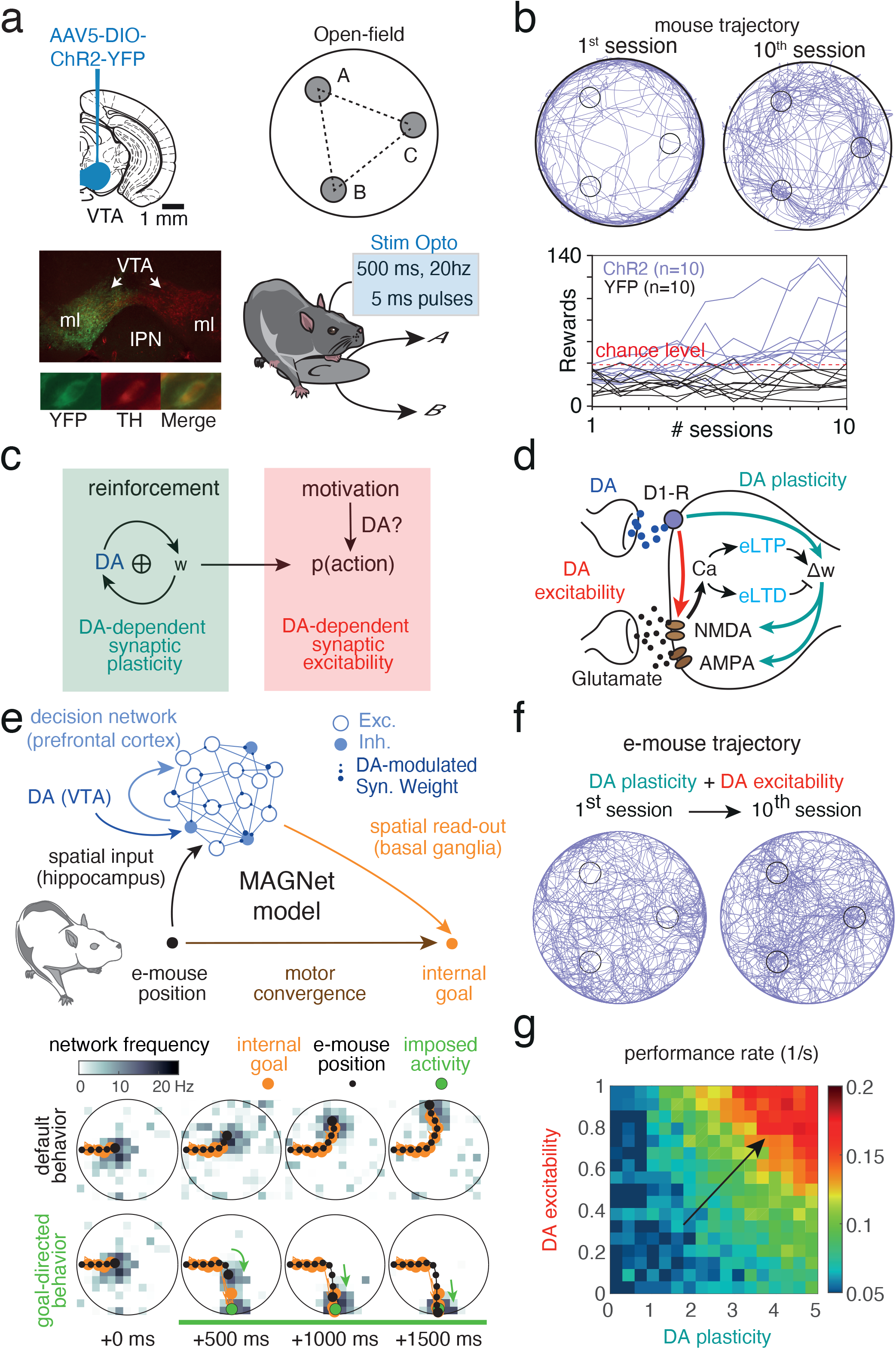
Increase in behavioral performance does not disentangle reinforcing and motivational roles of dopamine. a. Left, ChR2-YFP-expressing virus was injected in ventral tegmental area (VTA) TH-expressing dopamine (DA) neurons in DAT-Cre mice. Right, VTA photostimulation was delivered when mice were detected within one of three explicit locations (A, B, C) in the open field. Mice could not receive two consecutive photostimulations at the same location (e.g. C), so they alternated between locations (e.g. A or B after C). b. Top, trajectories (10 min) of one mouse expressing ChR2 in the VTA (purple) at the beginning (left) and at the end (right) of the learning sessions. Bottom, number of photostimulations against session number for Chr2-expressing (purple) and YFP-expressing (black) animals. c. DA reinforces synaptic weights used for decision-making, by increasing the probability actions that previously led to reward (green panel). Short-term DA effects on synaptic excitability may also increase the probability of upcoming actions (red panel), but the specificity of such a motivational role remains unclear. d. Modeling DA biophysics in the MAGNet model. At excitatory synapses, DA consolidates calcium-induced early eligibility traces (eLTD and eLTP) into long-term weight changes affecting AMPA and NMDA currents (DA-plasticity; green arrows). DA also instantaneously upregulates NMDA maximal conductances (DA-excitability; red arrow). e. Top, the decision architecture of the MAGNet model comprises a biophysical prefrontal (PFC) recurrent network with DA-modulated excitatory synapses, hippocampal position-encoding inputs (black), basal ganglia internal goal soft-max-decoding (orange) and motor convergence toward the internal goal (brown). Bottom, hippocampal inputs impose an activity bump (dark gray in maps). Under default behavior (upper maps), the e-mouse position (black dot) and internal goal (orange dot) are conjoined, such that goal-directed behavior is inoperant and navigation oriented toward the wall. Under goal-directed behavior (lower maps), the internal goal is decoded at a larger, distant, activity peak (artificially created here, green dot) such that the e-mouse converges to the goal (arrows). f. e-mouse trajectories during the 1st and 10th session of simulated protocol where DA-plasticity and DA-excitability operated online. g. Average number of rewards (performance rate) as a function of DA-plasticity (maximal Hebbian Assemblies weight) and DA-excitability (NMDA scaling factor).

In reinforcement learning (RL) theories, phasic DA signals reward prediction errors and teaches stimulus-action values by reinforcing weights linking sensory states to rewarded actions (*4*), consistent with experimental evidence that DA enables long-term synaptic plasticity in cortical/subcortical areas (*6*, *7*). However, DA also modulates effective synaptic efficacy by instantaneously potentiating N-methyl D-aspartate (NMDA) currents which are paramount in setting network dynamics (*5*, *33*), but overlooked in RL theories.

To dissect how these dual biophysical roles for DA in reinforcement (DA-plasticity) and motivation (DA-excitability) interact (Figure 1c) to account for decision-making in our optogenetic DA-conditioning task (Figure 1a-b), we developed the MAGNet model consisting of a distributed decision architecture assessing how an artificial mouse (e-mouse) navigates under DA regulation. Simulated phasic DA was delivered when the e-mouse crossed the rewarded locations, but also randomly during navigation to account for spontaneous DA occurring in mice (*14*, *17*) (see Methods). Notably, the model accounts for DA consolidation of spike-timing dependent plasticity (STDP) eligibility traces (light blue) into excitatory synaptic changes (DA-plasticity; green arrows), instantaneous DA NMDA upregulation (DA-excitability; red arrow), or a combination of the two (DA-plasticity-excitability; Figure 1d and Methods). These excitatory synapse models were embedded within a biophysical prefrontal (PFC) recurrent circuit model (Figure 1e, upper panel, blue) (*34*, *35*) (see Methods), with mixed reward-space neuronal selectivity (*35*–*37*). As place-reward association relies not only on the PFC, but also on a distributed architecture encompassing basal ganglia, thalamus, hippocampus and amygdala (*38*, *39*), we designed the MAGNet model as a distributed decision architecture (see Methods). The PFC network was organized topologically, neurons possessed preferred locations and received feed-forward inputs (black) encoding the mouse position, putatively from hippocampal place cells (*36*). In turn, an internal goal was decoded from PFC neurons activity and preferred positions using a soft-max selection rule representing basal ganglia operation (Figure 1e, orange). Finally, the e-mouse converged toward its internal goal with speed ballistics accounting for commands set by motor structures (Figure 1e, brown).

When network spiking was dominated by a bump of activity (gray squares, Figure 1e, lower panel, upper maps) arising from hippocampal inputs encoding e-mouse position (black dot), the internal goal (orange dot) and e-mouse position were confounded. Thus, behavior was not goal-directed and navigation was governed by default behavior toward and along arena walls (with some inroads into the arena; see Methods, Figure 1f, first session left). By contrast, when a larger activity bump was associated with a position distant from that of the e-mouse (as artificially introduced for illustration in Figure 1e lower map, green dot), the internal goal rapidly shifted to that position and navigation was dominated by goal-directed convergence (Figure 1e, green arrows).

As we observed with real mice (Figure 1b), e-mice learned three place–reward associations when navigating in the arena (Figure 1f). Moreover, while increasing DA-excitability alone was unable to trigger learning or affect behavioral performance, it substantially enhanced the effect of DA-plasticity on performance (Figure 1g). Hence, multiple synergistic combinations of increases in both DA-excitability and DA-plasticity could account for our experimental data, suggesting, in turn, that RL-type explanations of decision-making exclusively based on DA-plasticity may be incomplete.

In order to disentangle long-term (DA-plasticity) from online (DA-excitability) effects of DA signal on decision-making, more constrained experiments are necessary. We thus assessed the role of DA-plasticity and DA-excitability independently within a simpler version of the MAGNet model, with a single rewarded location at the arena center (Figure 2a). With DA-plasticity only (Figure 2b), simulated phasic DA delivered when the e-mouse crossed the rewarded location (which occurred by chance in naïve e-mice, see Methods) yielded longterm synaptic plastic modifications (top panels) that accumulated over trials (bottom). The resulting strongly connected Hebbian assembly encoded the place–reward association (right panel). By contrast, DA-excitability only transiently increased synaptic efficacy (<1s) in all the network, as a consequence of NMDA potentiation on a short timescale (Figure 2c).

**Figure 2.**
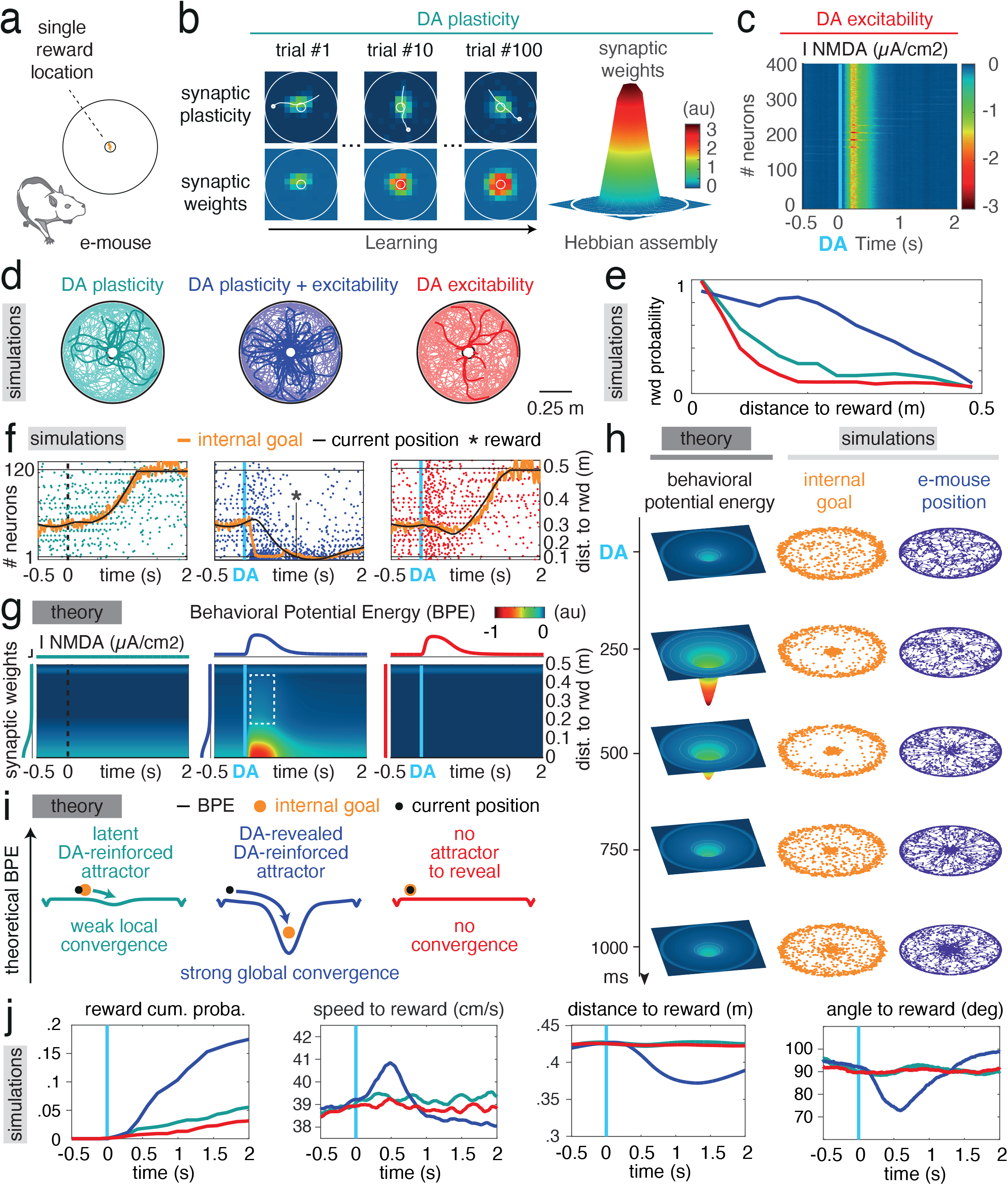
Dopamine builds and reveals latent network attractors encoding internal goals. a. Schematics of the single rewarded location arena. b. Under DA-plasticity alone, phasic DA delivered at the rewarded location yielded longterm synaptic changes (top panels) that accumulated (bottom), eventually shaping a Hebbian assembly encoding the place–reward association (right). c. Under DA-excitability alone, DA transiently increased synaptic efficacy in the whole network through NMDA potentiation. d. Superimposed example e-mouse trajectories in the DA-plasticity, DA-excitability and DA-plasticity+excitability conditions, from random positions and directions. Rewarded trajectories are in bold. The color code for conditions is used in panels e-j. e. Reward probability as a function of the initial distance of trajectories in d, for the three conditions. f. Example model dynamics (neural spiking sorted according to the distance to reward) in the three conditions. Under DA-plasticity+excitability, DA generated a massive neural co-activation at the Hebbian assembly, setting the internal goal at the rewarded location and e-mouse convergence toward it (reward). The Hebbian assembly was generally unexpressed under DA-plasticity, or absent under DA-excitability, forbidding goal-directed behavior. g. Theoretical behavioral potential energy (BPE) computed as a function of time and distance to reward under the three conditions. The rewarded location becomes a transient attractor of behavioral dynamics only under DA-plasticity. Faint blue strip at the top reflects the propensity to follow walls during default behavior. h. Theoretical BPE (illustrated in 2D), as well as internal goal and e-mouse position of example simulations, under DA-plasticity-excitability, during the first second following phasic DA. i. Schematics of attractorial dynamics in the MAGNet model. Theoretical BPE were computed at their maximal amplitude after phasic DA in the three conditions. Under DA-plasticity-excitability, navigation toward the reward arises from convergence toward the BPE minimum, which sets the internal goal. j. Ballistic model predictions indicate an increased cumulative reward probability under DA-plasticity-excitability only, due to an increase of speed to reward, as well as a specific approach to it (decrease of distance and angle to the reward).

When navigating in the arena, the e-mouse converged more toward the rewarded location if DA-plasticity and DA-excitability were considered simultaneously (Figure 2d, center, bold trajectories), rather than separately (Figure 2d, left and right). Moreover, convergence occurred from more distant positions in the DA-plasticity+excitability condition, whereas it was essentially local under either DA effect taken in isolation (Figure 2e). Indeed, in the DA-plasticity+excitability condition, modeling instantaneous NMDA potentiation (i.e. DA-excitability) had a larger, multiplicative effect on synapses already potentiated by DA-plasticity, resulting in a massive co-activation of neurons from the Hebbian assembly (Figure 2f). DA-plasticity+excitability thus set the internal goal (orange trajectory) on the learned reward location, attracting the e-mouse (black trajectory).

These biophysically-informed e-mouse simulations thus suggested a peculiar dynamical mechanism by which instantaneous DA-excitability reveals long-term DA-plasticity reinforcement and drives decision-making in mice. We developed a formal account of these complex systemic interactions (see Methods). In the MAGNet theory, analytically derived from the biophysical model, e-mice behavior could be reduced to one-dimensional dynamics and a behavioral potential energy (BPE) could be determined as

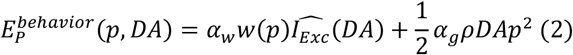

with *p* the e-mouse position 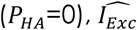 the weight-normalized excitatory current, *w* the sum of DA-reinforced synaptic weight, and *α_w_*, *α_g_* and *ρ* constants (see Methods). This theoretical expression reveals that convergence to the rewarded location was dictated both by (1) strong, local attractor dynamics, where the progressive increase in synaptic weights nearby the Hebbian assembly works to destabilize and attract non-goal-directed neural activity (Figure 2g, center, red spot), and (2) weaker, global attractor dynamics due to focalization of the internal goal at the Hebbian assembly (dashed box). Both of these terms required instantaneous DA-excitability action on a previously DA-plasticity-reinforced Hebbian assembly, as they were negligible when either DA-plasticity or DA-excitability was absent (Figure 2g left, right). Altogether, under the DA-plasticity+excitability condition, phasic DA signaling induced the transient unfolding of a large and deep BPE basin of attraction (Figure 2h, left), subsequent focalization of the internal goal (Figure 2h center) and, ultimately, e-mouse convergence to the rewarded location (Figure 2h right).

According to MAGNet model simulations and theory, DA-plasticity generates *latent* attractors only allowing weak local convergence of internal goal and e-mouse positions (Figure 2i, left). DA-excitability reveals these latent attractors, by amplifying their depth and width, resulting in strong global convergence (Figure 2i, center), which is impossible without previous learning (Figure 2i, right). The model allowed us to make ballistic predictions to describe reward-seeking behavior in the e-mouse. Specifically, under DA-plasticity+excitability, the cumulative probability of convergence to the rewarded location grew faster compared to other conditions (Figure 2j, first panel). This effect arose from both an energization of e-mice, with increased speed (second panel) and reorientation, as well as a decrease in the e-mouse’s distance to the reward (third panel), due to reorientation of their approach angle toward the reward location (last panel).

To test these predictions, we restricted VTA photostimulation in mice to directly evoke DA-excitability, and used electrical stimulation instead to promote DA-plasticity processes during learning. Mice were implanted with an electrode in the medial forebrain bundle (MFB), as electrical stimulation of the MFB has long been considered as a potent substrate for brain stimulation reward (*40*), and an optic fiber in the VTA (four weeks after viral injection of ChR2, as in Figure 1, Figure 3a). We used MFB stimulation to build the reward-place association in the circular open-field context (Figure 3b), where the center location was paired with twenty 0.5 ms electrical pulses at 100 Hz, and mice were required to leave the location before being stimulated again upon reentry (*30*, *31*). This led to strong reinforcement of the central place preference (F(9)=5.57, p = 0, Figure 3c, Supplementary Figure 2), so that current intensity was adjusted to achieve a moderate visit rate.

**Figure 3.**
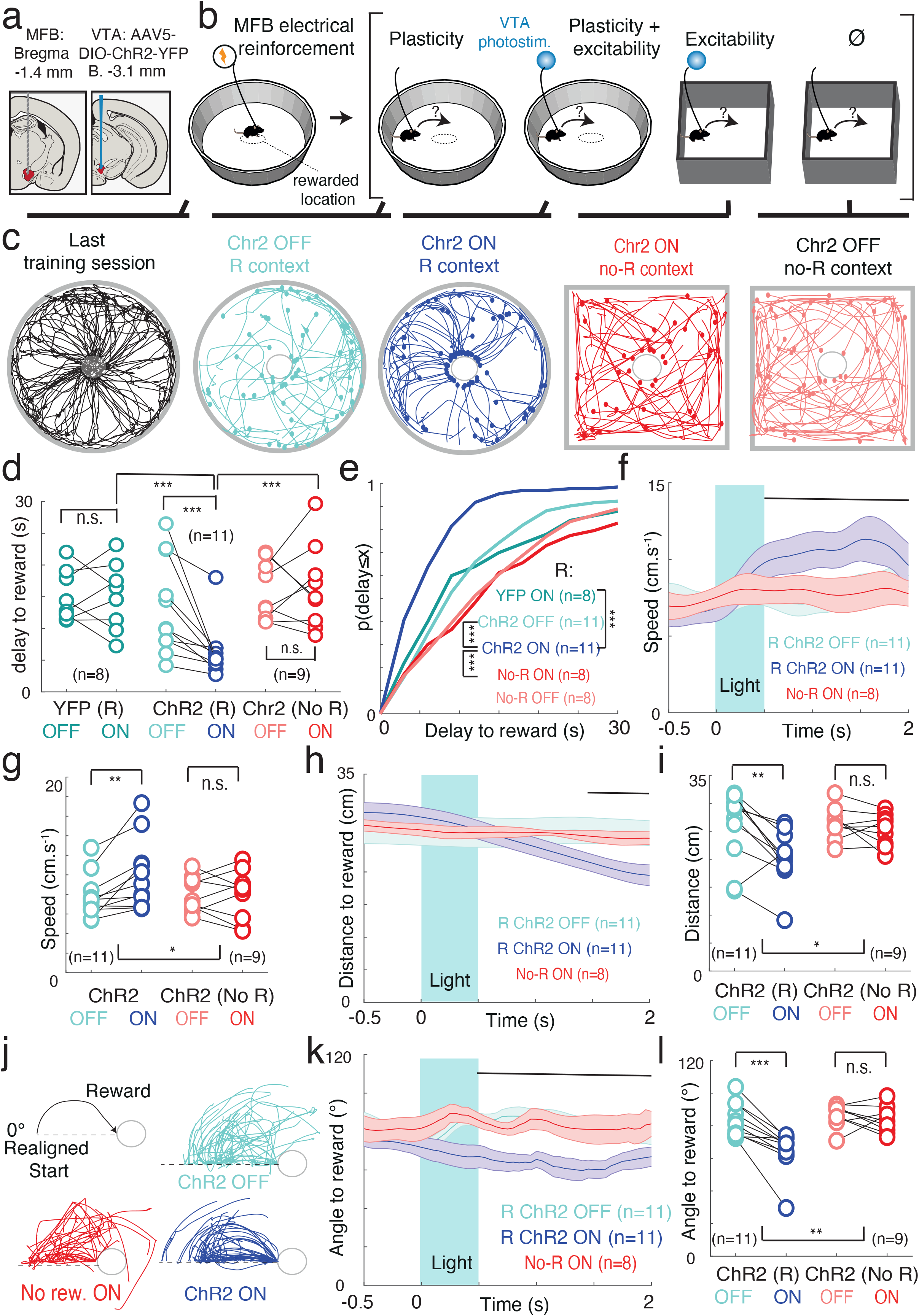
Testing the prediction that VTA photostimulation-induced movements are goal-specific and context-dependent. a. Schematics of electrode implantation in the medial forebrain bundle (MFB) and injection of the ChR2-YFP-expressing virus and fiber implantation in ventral tegmental area (VTA). b. Experimental test of the model predictions. A location is rewarded by MFB electrical stimulation (left). Then (inside the brackets), VTA photostimulation is provided in the context where reinforcement occurred (plasticity + excitability) and compared to a “plasticity only” conditions (MFB+Chr2 animals without VTA photostimulation), an “excitability only” condition (VTA photostimulation in another context where no location had been rewarded), and “null” condition (no photostimulation and no reward context). c: From left to right : example trajectories at the end of the MFB conditioning sessions, and post-photostimulation bouts of trajectories in the different conditions described in c. Differences between photostimulation-rewarded location delays for YFP (reward context); ChR2-expressing (reward context) and ChR2-expressing (no reward context) animals d. Cumulative distribution of the photostimulation-rewarded location delays in YFP (ON light in reward context); ChR2-expressing (ON and OFF light in reward context) and ChR2-expressing (ON and OFF light, no reward context) animals. e-l. Speed (d), distance to the rewarded location (f), and angle between the animal and the rewarded location (i) around VTA photostimulation for ChR2-expressing animals when ON light in reward context (purple), OFF light in reward context (light blue) and ON light in no reward context (red). Average difference in speed (e), distance to the rewarded location (g), and angle between the animal and the rewarded location (j) between ON and OFF light conditions, in reward (“Chr2”) and no reward (“Chr2 No R”) contexts. (h) shows the computation of angle between the animal and the rewarded location, based on the same trajectories as in a, realigned to the same line relative to the rewarded location, showing straight trajectories for animals when ON light in reward context (purple), and indirect trajectories when OFF light in reward context (light blue) or ON light in no reward context (red).

Once the association was learned, we used VTA photostimulation (which mice had never encountered before, controlling that the LED was not used as a cue with YFP-transduced mice) with similar parameters as we used previously for DA-plasticity (reinforcement) (Figure 1). To test instead for the motivational effect of phasic VTA DA signaling, we provided brief photostimulations when mice were either (see Figure 3b for the different contexts and Figure 3c for illustrative examples of trajectories) in the same open-field, but away from the reinforced position (reward context, R), or in a different context (square open-field), where no location had been associated with a MFB reward (no-reward context, no-R). This allowed for experimentally dissecting the role of the two different DA effects identified in the MAGNet model in the decision-making behavior of real mice. In this real-world test, the DA-plasticity condition (i.e. with only baseline DA-levels during ongoing decisions) consisted of mice tested in R context, but not receiving effective VTA photostimulations (YFP ON or Chr2 OFF + reward context), which only expressed previous reinforcement and default behavior. Mice in the DA-plasticity-excitability condition (i.e. with increased level of DA during ongoing decisions) received VTA photostimulations while tested in the R context (Chr2 ON + reward context), which resulted in increased phasic DA signaling during decision-making. Finally, DA-excitability mice received online VTA photostimulations in a no-R context (Chr2 ON + no reward context) in the absence of previous plasticity in such a no-R context.

In ChR2-expressing mice tested in the R context, VTA photostimulation decreased the delay to the reward location compared to control times (Figure 3d, R/ChR2 ON versus OFF paired t-test: T_(10)_= −3.58, p=0.05, and Figure 3e, KS test on all trials from all mice: p=1.10^−8^). This effect was neither observed in YFP-expressing animals (R/YFP ON versus OFF, paired t-test: T_(7)_= −0.07, p=0.94, KS test on all trials from all mice: p=0.23). By contrast, VTA photostimulation did not reduce the delay to the center location in ChR2-transduced animals in the no-R context (No-R/ChR2 ON versus OFF, paired t-test: T_(8)_= 0.32, p=0.76, KS test on all trials from all mice: p=0.81). Hence, a decrease in the latency to visit the rewarded location was only observed in the experimental equivalent of the DA-plasticity+excitability condition, as predicted by the MAGNet model (Figure 2j, first panel).

We next investigated whether this reduced delay following VTA photostimulation reflected an increase in speed (Figure 3f), i.e. an energizing effect (*3*, *17*) rather than an increase in the overall pace (frequency) of behavior (*41*). VTA photostimulation in the reward context resulted in an increase of animal speed (Figure 3g, R/ChR2 ON versus OFF paired t-test on speed after stimulation: T_(10)_=3.46, p=0.006, see Methods), which was neither observed in YFP controls (T_(7)_=−0.44, p=0.67, Supplementary Figure 2), nor in ChR2 animals in the no-R context (T_(8)_=−0.17, p=0.87). Online manipulation of VTA DA signaling during decision-making behavior thus affected the speed of action (rather than just its pace), but only in the context in which a place–reward association had already been made, again consistent with the MAGNet model prediction (Figure 2j second panel). Hence, VTA DA signaling only exerted an energization effect in the reward context, which is incompatible with decision threshold models predicting context-independent speed increases (*3*, *13*). The MAGNet theory, based on attractor dynamics, also predicts, contrary to RL or decision-threshold models, that the increase in speed following DA stimulation would be directed towards the reinforced location. We thus assessed whether online VTA DA signaling also affected mice directional behavior. First, the distance between ChR2-transduced animals and the central location (Figure 3h) decreased upon VTA DA photostimulation in the R context (Figure 3i, T_(10)_=−3.68, p=0.004) but not in YFP animals (T_(7)_=−0.92, p=0.39, Supplementary Figure 2), nor in ChR2 animals in the no-R context (T_(8)_=−1.17, p=0.27). Second, the accumulated sum of successive angles between the animal and the goal (error angle, Figure 3j) decreased following stimulation in ChR2-expressing animals in the R context (Figure 3k,l; paired t-test stimulation vs control: T_(10)_=−5.32, *p* = 3.10^−4^) indicating more direct trajectories to the reward, rather than faster trajectories in any direction. This was neither the case in YFP-expressing mice (T_(7)_=−0.47, p=0.66, Supplementary Figure 2), nor in ChR2 animals in the no-R context (T_(8)_=−0.89, p=0.40).

Hence, the increase in animal speed following optical stimulation of VTA DA neurons was directed toward the central location, consistent with the MAGNet model’s expectations that phasic, online DA signaling would exert a goal-directed energizing effect. DA signaling only attracted the animals toward the center location in a context in which this location had been previously associated with MFB stimulation reward, suggesting that instantaneous DA-excitability (motivation) acts in a content-specific and context-dependent manner to retrieve the goal learned under DA-plasticity (reinforcement).

## DISCUSSION

The biophysically-informed MAGNet dynamical theory of DA actions interprets goal-directed actions as a two-step process: neural assemblies representing a potential goal are learned through DA-regulated synaptic plasticity, but not automatically expressed, i.e. they are latent in terms of behavior (*42*–*44*). Then, phasic DA signaling has the ability to make these attractors accessible from remote starting conditions, by widening and deepening their basins of attraction. We validated the MAGNet theory experimentally using optogenetics, showing that online phasic DA signaling orients the animal toward rewarded locations and energizes specific, context-dependent actions previously entrained by phasic DA-induced plasticity. We thus propose that phasic DA signaling biases how ongoing decisions are being made by controlling the landscape of potential behaviors on a fast timescale.

## AUTHORS CONTRIBUTIONS

JN BD PF designed the study. BD JN MS designed the model, and performed the simulations with FA. JN BH performed the behavioral experiments. SM EV JN performed viral injections and histology. JN PF analyzed the behavioral data. AM LT contributed tools/reagents. JN MS PF BD wrote the article.

## ACKNOWLEDGEMENTS

We thank Deniz Dalkara and the viral core facility at the Vision Institute (Paris) for the AAV vectors. We thank Lauren Reynolds and Benoît Girard for careful reading of the manuscript.

This work was supported by Centre national de la recherche scientifique (CNRS UMR 8249), Fondation pour la recherche médicale (FRM DEQ2013326488 to PF, FRM FDT201904008060 to SM), French National Cancer Institute Grant TABAC-16-022, French state funds managed by Agence nationale de la recherche (ANR-16 Nicostress to PF, ANR-20 NICADO),

## SUPPLEMENTARY DISCUSSION

### Biophysical network modeling of behavior

Contrary to reinforcement learning models that focus on the phenomenology of behavior rather than on biological implementation, the MAGNet model constitutes an attempt to root a dynamical theory on biophysical and biochemical properties, relying on three key features.

First, we considered a recurrent network, because it links attractor dynamics to elemental computations of decisions (*1*). Even if inspired by the cortical stage of decisionmaking (*2*, *3*), the MAGNet model does not exclude other parts of the mesocorticolimbic loop from the decision process (*4*, *5*). In particular, striatal dopamine is needed for approaching rewards (*6*, *7*), and our theoretical proposal that online dopamine affects the behavioral potential energy is based on the convergence of the whole decision architecture toward an attractor, encompassing the striatal stage. In decision-making models based on basal ganglia circuits, navigation toward goals can be learned through reinforcement-learning of synapses between space- (e.g. hippocampal) and action-coding (striatal) neurons (*8*). Other models have proposed a link between action selection and action intensity (*9*), accounting for the role of basal ganglia in energizing behaviors (*10*). DA regulation on both synaptic plasticity and excitability could thus result in multiplicative effects of DA (*11*) on action selection and energization in a striatal model combining these different features. Such a combined model remains to be achieved. Nevertheless, if the online DA effect in striatal networks is to increase the gain of action selection (*12*), then online DA may favor space-action sequences leading to reinforced locations. However, navigation models of basal ganglia are not based on attractors at the level of neural network dynamics. Instead, convergence toward a goal corresponds to the animal progressively following gradients of space-action values (*8*), analogous to the local convergence along synaptic weight gradients in our model. Hence, gathering these different striatal models together could hypothetically account for the deepening of the goal’s basin of attraction, although this remains to be shown. However, it seems more difficult for such neurodynamical systems to account for the widening of the goal’s basin of attraction, which requires a distant signal focalizing the dynamics of the internal goal, from any initial condition. Altogether, deciphering the respective effects of dopamine on corticostriatal NMDAR and on the intrinsic excitability of medium spiny neurons compared to NMDAR from recurrent connections would refine the link between MAGNet model’s predictions and the neurobiology literature. Finally, DA is likely to also affect online the amygdala, thalamus and hippocampus, as well as the connections between these structures and the cortex and basal ganglia (*4*, *13*), such full-scale modeling being out of scope. We thus lumped some of the decision processes into simple (e.g. spatial coding as a topographical feed-forward excitatory input) or phenomenological descriptions (e.g., motor convergence as linear and angular ballistics commands).

The second important feature of the MAGNet model concerns plasticity pathways implementing eligibility traces with synaptic tags. We followed the recent literature describing two distinct eligibility traces for LTP and LTD (*14*), but this separation leaves holes in the implementation by intracellular pathways. Indeed, early LTP and LTD are believed to depend on CaMKII and calcineurin, respectively, while in the present model a different couple of kinase and a phosphatase is needed for LTP and LTD. This may be implemented by compartmentalization via synaptic scaffolds linking different forms of CaMKII with different phosphatases (*15*). Likewise, downstream decoding of early LTP/D may be achieved by ERK and CREB (*16, 17*), although these steps may not be specific to glutamate receptor upregulation. A more refined model would require to include other DA regulations (*13*, *18*), such as intrinsic and structural plasticity. In the MAGNet model, dopamine is key to transform eligibility traces into effective plasticity, but other neuromodulators such as noradrenaline (NE), serotonin (5HT) and acetylcholine (ACh) seem to exert differential effects on the read-out of LTP and LTD (*14*). Linking the behavioral events that triggered these neuromodulators, together with the precise form of eligibility mechanism they implement, would enrich our comprehension of reward (or other outcomes)-gated plasticity.

The third key feature is the online modulation exerted by DA. Here, we focused on NMDA modulation, whereas DA can affect a vast diversity of receptors and ionic channels depending on the structure and the subtype of DA receptors (*13*, *19*). Similarly, we mainly modeled D1R effects, but D2R may not be as antagonistic as previously believed: D1R and D2R are actually synergistic when considering the cAMP-PKA pathway we considered (*13*). Even the regulation of intrinsic excitability of medium spiny neurons is more complex than D1R-mediated increases in excitability and D2R-mediated decreases: D2R may exert destabilizing influences (rather than inhibitory) that promote or oppose D1R effects depending on down or up-states, respectively (*13*). These interactions hint at complementary roles in our dynamical framework, that we discuss below.

### Relations to other theories of dopamine function

Reinforcement learning theories do not assign any effect to dopamine during ongoing behavior, once the value of actions has been learned through DA modulation of plasticity (*20*, *21*). In alternative views to RL, dopamine has been suggested to exert either directional effects, i.e. stimulus-driven dopamine release directs the behavior toward the cue (*22*, *23*) or activational effects, i.e. dopamine increases the probability and vigor of any motor behavior (*10*, *24*). Both views explain some of the vast literature on phasic dopamine. DA nuclei do not have enough encoding capacity and DA projections are not selective enough (*10*, *24*) to precisely represent the goal toward which the animal should be directed. As such, in the directional account of dopamine, DA is proposed to add incentive motivation or salience to the cue being currently processed, promoting approach through yet-unknown mechanisms. The DA-associated cue is described in incentive-salience accounts as becoming “magnetic” (*22*, *23*), which is exactly what is expected in the MAGNet model for a state suddenly attracting the decision network’s dynamics. However, actions that are not cue-driven but self-generated rely on internal representations, in which case the role of DA in incentive-salience is less specified. Our proposal is based on contextual decisions, in which animals rely on learned internal representations to approach reward. It reinterprets incentive motivation as making attractors (stable steady states) representing potential (sensory or internal) goals accessible to neural dynamics. However, DA does not support a purely directional role in our theory. Indeed, although DA promotes convergence toward distant rewarded goals in a given context, the chosen goal is not itself specified by DA neurons. Rather, choice results both from the current animal position and the BPE landscape built by previous reward history: animal will converge toward the goal owning the basin of attraction in which it lies at the time of phasic DA.

Activational accounts assign a general role to phasic dopamine in gating decisions (increasing DA would render all decisions more probable) and energizing actions. Incentive motivation models, in which decisions are sequentially evaluated, i.e. accepted or not based on the intensity of phasic DA (*25*), would predict an undirected increase in the probability of every action following VTA photostimulation, in opposition to our experimental data showing a reduced angle to reward, and an absence of DA effects outside the reward context. Furthermore, we show that speed profiles, not just latency or average speed, are affected by phasic DA, which go beyond the scope of discrete-time models (*25*). Phasic DA has also been suggested to move the threshold for decisions in drift-diffusion models (*10*, *24*) predicting context-independent increase in undirected actions, which is also inconsistent with our observations on context-dependent directed energization of actions. Widening the basin of attraction in the present model naturally increases both the likelihood, directness and speed of actions in a reward context-dependent fashion. Finally, in time-processing accounts, dopamine affects the sense of time: under high DA time goes fast, while under low DA time is felt as slower (*26*). We interpret this as the speeding up of neuronal and behavioral dynamics upon attractor unveiling. In the context of working memory, tonic levels of prefrontal DA have been related to the gating and maintenance of persistent activity encoding a goal (*19*, *27*). In this account, D2R favors stimulus-driven transitions toward another goal by rendering attractors more shallow, while working memory of the current goal is stabilized by D1R-mediated deepening of its basin of attraction. This view differs from ours, in which phasic DA activates D1R to both widen and deepen the basin of attraction, setting a goal based on an internal memory. DA roles in decision-making and working memory are not necessarily opposite, as DA may achieve a “double duty” in cognitive motivation (*28*) by widening (to promote the decision) and deepening (to stabilize its working memory) basins of attraction. Here, the experimental test considered the physical space of the open-field as the task space. However, the conceptual consequences of the MAGNet model extend to non-physical spaces. Indeed, dopamine is needed for approaching rewards when animals are far in terms of either physical or task space (*6*, *28*). Hence, our MAGNet theory translates the “flexible approach” (*6*) hypotheses into a neurodynamical process: online DA is needed to approach reward in non-habitual situations, before the animal engages toward its goal, or when there is a motor cost and that DA is needed to travel some distance (in either physical or task spaces) to retrieve specifically DA-associated goals.

In the experimental literature, exogenous stimulation of phasic DA has provided conflicting results, with context-independent (*29*) and context-dependent (*30*) movement following SNc/dorsal striatum stimulation. Stimulation of VTA DA neurons exerts either context-dependent effects (*31*), or fails to affect online behavior (*32*). It has notably been advocated that phasic DA would only affect online behavior if animals are preparing to move (*33*). Our theory reconciles these conflicting results: when the animal is head-fixed, already close to a rewarded state (*32*), VTA DA is unneeded and its stimulation does not change behavior (i.e. as when the e-mouse is already at its internal goal in the model). In contrast, for situations in which animals must travel in a physical or task space, DA increases the likelihood and speed of convergence toward a goal (*30*, *31*). For the same reason, our framework also explains why no dopamine is needed for no-go conditioning (*34*), as such a case would correspond, in the present model, to the goal state being the rest state. Furthermore, the dichotomy between SNc and VTA may be based on the type of attractor these nuclei affect. VTA could build and express high-level goals (deep and substantially separated wells in the energy landscape), and SNc low-level, context-independent subgoals (i.e. a given motoric action) corresponding to multiple nearby attractors, explaining context-independent locomotion upon SNc stimulation (*29*).

### Latent attractor as a new dynamical framework distinguishing learning from performance

It is difficult to assign biological meaning to parameters of the phenomenological models discussed above (i.e. reinforcement learning and their generalizations). Biophysically-based neurodynamical models can help bridge description levels in neuroscience, i.e. build on cellular and network mechanisms to account for behavior (*35*). Neurodynamical theories ascribe the property to self-sustain stable neural activity - also known as attractor states - to recurrent networks underlying decisions, such as the prefrontal cortex (*35*). In the context of decision-making, attractors in the neural state space would represent external and/or internal information encoding potential decisions (*36*). Hence, decision-making in recurrent networks depends on transitions between the spontaneous activity state (i.e. corresponding to no decision), and decision attractors (*37*). Such transition may be achieved by a stimulus cue acting as an input to the network, which destabilizes the spontaneous state and transforms it into one of the decision states toward which dynamics will slowly evolve (*1*). Alternatively, the spontaneous state and decision states may coexist as several stable attractors (multistability), with fluctuations in neural activity (noise) driving the transition toward decision states (*37*). However, neither models based on noise nor on stimuli can explain self-generated, goal-directed actions in which the decision is taken based on internal representations (*38*).

Our proposal solves this issue by proposing that DA reveals decision attractors (hereby destabilizing the rest state) which would not express otherwise. Such revealing is allowed in the MAGNet model by considering two distinct yet linked spaces, the cognitive space and the behavioral space, which would coincide in most models. Here, a circular causality links mouse navigation in the behavioral space and internal network representations in the cognitive space, resulting from feed-forward inputs encoding the mouse position, downstream decoding of neural activity into an internal goal, as well as weighted recurrent connectivity learned through past dopaminergic reinforcement (DA-plasticity), and online motivational dopaminergic modulation of effective synaptic efficacies (DA-excitability). Hence, we do not consider the local convergence of neural activity toward an attractor solely in neural state space, but rather the convergence in the merged neural and behavioral space (i.e. their cartesian product). This allows DA to exert a distant, discontinuous role (the internal goal is instantaneously set at a distant position) which widens the decision’s basin of attraction, at odds with local effects on attractor stability (deepening).

This new principle also provides a simple solution to the common critique addressed to attractor models that real neural activity is never actually stationary, but transient. Indeed, in standard, Hopfield-like models, decision-making comes to a standstill once the activity closes in at the attractor (a steady-state with only attracting dimensions). More refined models consider saddles (*39*) or attractor ruins (*36*), i.e. partially stable attractors, with attracting, stable directions co-existing with unstable directions), allowing dynamics to eventually escape and converge to another attractor (*36*, *39*). This requires specific mechanisms, either synaptic inhibition designed to repel the neural dynamics from the attractor (*39*) or neuronal fatigue ensuring the attractor to be only transient once activated (*40*). Contrary to these models, the decision attractor simply vanishes in the MAGNet model, once the excitability effect of phasic DA decays due to recapture. Hence, both the entry into, as well as the exit from, the decision attractor are controlled by an internal operation (i.e. a motivational state implemented by phasic DA) in our theory. Such internal control also effectively decouples the neural dynamics from synaptic changes, which is key to account for goal-directed actions. Usually, reward-dependent synaptic plasticity directly leads to a change in models’ neural dynamics, yielding behavioral adaptation (i.e. change in the frequency of behavior). However, animals do not always express learning as behavioral changes. Instead, some forms of learning are latent (*38*, *41*, *42*). For instance, a sated animal may learn to navigate a labyrinth containing a food source without increasing the visits to the food source, and, upon food deprivation, display a change in its behavior (i.e. going to the food source). The MAGNet model accounts for such latent learning by dopamine-modulated synaptic plasticity only building latent attractors that do not necessarily affect neural dynamics. We thus provide a neurodynamical account of how motivation, implemented by phasic DA, is needed to express the memory of previous, latent, reward learning.

## METHODS

### Animals

Experiments were performed on DAT^iCRE^ female (n=26) and male (n=21) mice, from 8 to 16 weeks old, weighing 25-35 grams. Mice were housed in cages in an animal facility where the temperature (21+/- 1°C) and a 12h light/dark cycle were automatically monitored with food and water available ad libitum. DAT^iCRE^ mice (*43*) were kindly provided by Ludovic Tricoire and genotyped by PCR as described previously (*44*). All experiments were performed in accordance with the recommendations for animal experiments issued by the European Commission directives 219/1990, 220/1990 and 2010/63, approved by Sorbonne University, and n° 014378.01 supervised by the CEEA - 005.

### Virus production

AAV vectors were produced as previously described (*45*) using the co-transfection method, and purified by iodixanol gradient ultracentrifugation (*46*). AAV vector stocks were titrated by quantitative PCR (qPCR) (*47*) using SYBR Green (Thermo Fischer Scientific).

### Virus injections

Mice were anesthetized with a gas mixture of oxygen (1 L/min) and 1-3 % of isoflurane (Piramal Healthcare, UK), then placed into a stereotaxic frame (Kopf Instruments, CA, USA). After the administration of an analgesic (Buprecare 0,1 mL at 0,015 mg/L) and of a local anesthetic (Lurocain, 0.1 mL at 0.67 mg/kg), a median incision revealed the skull which was drilled at the level of the VTA. Mice were then injected unilaterally in the VTA (1 μL, coordinates from bregma: AP −3.1 mm; ML ±0.5 mm; DV −4.5 mm from the skull) with an adeno-associated virus (AAV5.Ef1a.DIO.ChR2.YFP 6.89e13 vg/mL or AAV5.Ef1a.DIO.YFP 9.10e13 vg/mL). A double-floxed inverse open reading frame (DIO) allowed to restrain the expression of ChR2 to VTA dopaminergic neurons. After stitching and administration of a dermal antiseptic, mice were then placed back in their home-cage and had 14 days to recover from surgery.

### Fiber and electrode implantations

Two weeks after virus injection, mice were anesthetized as above. After the administration of the analgesic and local anesthetic, skin was incised, the skull was drilled at the level of the VTA. An optical fiber (200 μm core, NA=0.39, Thor Labs) coupled to a ferule (1.25 mm) was implanted just above the VTA ipsilateral to the viral injection (coordinates from bregma: AP −3.1 mm, ML ±0.5 mm, DV 4.4 mm), and fixed to the skull with dental cement (SuperBond, Sun Medical).

For dual implantation experiments, the skull was also drilled at the level of the Median Forebrain Bundle (MFB). A bipolar stimulating electrode was then implanted unilaterally (ipsilateral to the optic fiber in the VTA) in the brain (stereotaxic coordinates from bregma according to Paxinos atlas: AP −1.4 mm, ML ±1.2 mm, DV −4.8 mm from the brain).

After stitching and administration of a dermal antiseptic, mice were then placed back in their home-cage and had 14 days to recover from surgery. The behavioral task began 4 weeks after virus injection to allow the transgene to be expressed in the target dopamine cells.

### Ex vivo patch-clamp recordings of VTA DA neurons

To verify the functional expression of the excitatory opsin ChR2, 8-12 week-old male and female DATiCRE mice were injected with the ChR2-expressing virus as described above. 4 weeks after infection, mice were deeply anesthetized with an intraperitoneal (IP) injection of a mix of ketamine/xylazine. Coronal midbrain sections (250 μm) were sliced using a Compresstome (VF-200; Precisionary Instruments) after intracardial perfusion of cold (4°C) sucrose-based artificial cerebrospinal fluid (SB-aCSF) containing (in mM): 125 NaCl, 2.5 KCl, 1.25 NaH_2_PO_4_, 5.9 MgCl_2_, 26 NaHCO_3_, 25 Sucrose, 2.5 Glucose, 1 Kynurenate (pH 7.2, 325 mOsm). After 10-60 min at 35°C for recovery, slices were transferred into oxygenated aCSF containing (in mM): 125 NaCl, 2.5 KCl, 1.25 NaH_2_PO_4_, 2 CaCl_2_, 1 MgCl_2_, 26 NaHCO_3_, 15 Sucrose, 10 Glucose (pH 7.2, 325 mOsm) at room temperature for the rest of the day and individually transferred to a recording chamber continuously perfused at 2 ml/min with oxygenated aCSF. Patch pipettes (4–8 MΩ) were pulled from thin wall borosilicate glass (G150TF-3, Warner Instruments) using a micropipette puller (P-87, Sutter Instruments, Novato, CA) and filled with a potassium gluconate (KGlu)-based intrapipette solution containing (in mM): 116 K-gluconate, 10-20 HEPES, 0.5 EGTA, 6 KCl, 2 NaCl, 4 ATP, 0.3 GTP and 2 mg/mL biocytin (pH adjusted to 7.2). Transfected VTA DA cells were visualized using an upright microscope coupled with a Dodt contrast lens and illuminated with a white light source (Scientifica). A 460 nm LED (Cooled) was used both for visualizing YFP-positive cells (using a bandpass filter cube) and for optical stimulation through the microscope (with same parameters used for behavioral experiments: ten 5-ms pulses at 20Hz). Whole-cell recordings were performed using a patch-clamp amplifier (Axoclamp 200B, Molecular Devices) connected to a Digidata (1550 LowNoise acquisition system, Molecular Devices). Signals were low-pass filtered (Bessel, 2 kHz) and collected at 10 kHz using the data acquisition software pClamp 10.5 (Molecular Devices). All the electrophysiological recordings were extracted using Clampfit (Molecular Devices) and analyzed with R.

### Behavior acquisition and conditioning procedures

Experiments were performed using a video camera connected to a video-track system, out of sight of the experimenter. A home-made software (Labview National instrument) tracked the animal, recorded its trajectory (20 frames per s) for 10 min and sent TTL pulses to the electrical stimulator or LED device when appropriate.

Conditioning procedure with VTA DA photostimulation: three explicit square locations, marked on the floor, were placed in a circular open-field (67 cm diameter), forming an equilateral triangle (side = 35 cm). Each time a mouse was detected (by its centroid) in the area of one of the rewarding locations (area radius = 3 cm), a 500-ms train of ten 5-ms pulses at 20 Hz was delivered to the LED device. An ultra-high-power LED (470 nm, Prizmatix) coupled to a patch cord (500 μm core, NA=0.5, Prizmatix) plugged onto the ferrule was used for optical stimulation (output intensity of 10 mW). Animals could not receive two consecutive stimulations in the same location.

Conditioning procedure with MFB electrical stimulation: only one explicit location was marked on the floor, at the center of the open-field. Each time a mouse centroid was detected in the area (radius = 5 cm) of the location, a 200-ms train of twenty 0.5-ms biphasic square waves pulsed at 100 Hz was delivered to the electrical stimulator. Mice were required to leave the location (i.e. to be detected at least 10 cm from the central point) for the stimulation to be made available again. The training consisted of a block of 5 daily sessions of 10 min at 80 μA, followed by 5 daily sessions of 10 min in which ICSS intensity was adjusted (in a range of 20-200 μA) so that mice visited the central location between 20 and 50 times at the end of the training.

Test sessions with VTA DA photostimulation: after the end of the MFB electrical conditioning procedure, the optical stimulation patch cord was plugged onto the ferrule during at least one OFF day (maximum = 5) to habituate the animals, until the criterion (between 20 and 50 locations visits in 10 min) was reached again. On ON test days, photostimulation was delivered by the experimenter when the animal was outside of the reinforced location (at least 10 cm from the central point). When the experimenter clicked to stimulate, it had a 50% probability to deliver an actual TTL pulse leading to photostimulation, otherwise this time point was recorded as a control. In the square open-field test, occurring after the test session in the circular openfield, the procedure was the same, except that it took place in square open-field (side = 70 cm) without any mark on the center.

### Behavioral analyses and statistics

Stimulation-reward duration was computed as the time between the start of the photostimulation (or of the control time) and the first detection of the animal in the central location. Durations greater than 60s were excluded from the analysis for the sake of representations, but did not affect the statistical significance of the tests. Cumulative distributions of durations were computed by pooling stimulation-reward and control time-reward from all animals in one condition (e.g. ChR2 or YFP), with a 3-s time bin. Average delays to rewards were also computed for each animal. For all groups of mice, the trajectory was smoothed using a triangular filter before computing the instantaneous speed, which corresponds to the distance traveled by the animal between two video frames (every 50 ms) as a function of time. Mean acceleration following stimulation was taken as the time derivative of speed during the first second after stimulation. Angles to reward were computed as the angles between each successive position of the animal relative to the initial angle (at photostimulation or at control time). Angle error was taken as the mean of || *∑e^iθ^* || where *θ* are the successive angles to reward.

All statistical analyses were computed using Matlab with custom programs. Results were plotted as a mean ± s.e.m. The total number (n) of observations in each group and the statistics used are indicated in figure legends. Classical comparisons between means were performed using parametric tests (Student’s T-test, or ANOVA for comparing more than two groups) when parameters followed a normal distribution (Shapiro test P>0.05), and non-parametric tests (here, Wilcoxon or Mann-Whitney) when the distribution was skewed. Repeated-measure ANOVAs were used for longitudinal measures. Multiple comparisons were Bonferroni corrected.

### Immunochemistry

After euthanasia, brains were rapidly removed and fixed in 4% paraformaldehyde (PFA). After a period of at least three days of fixation at 4°C, serial 60-μm sections were cut with a vibratome (Leica). Immunostaining experiments were performed as follows: VTA brain sections were incubated for 1 hour at 4°C in a blocking solution of phosphate-buffered saline (PBS) containing 3% bovine serum albumin (BSA, Sigma; A4503) (vol/vol) and 0.2% Triton X-100 (vol/vol), and then incubated overnight at 4 °C with a mouse anti-tyrosine hydroxylase antibody (anti-TH, Sigma, T1299) at 1:500 dilution, in PBS containing 1.5% BSA and 0.2% Triton X-100. The following day, sections were rinsed with PBS, and then incubated for 3 hours at 22-25 °C with Cy3-conjugated anti-mouse and secondary antibodies (Jackson ImmunoResearch, 715-165-150) at 1:500 in a solution of 1.5% BSA in PBS, respectively. After three rinses in PBS, slices were wet-mounted using Prolong Gold Antifade Reagent (Invitrogen, P36930). Microscopy was carried out with a fluorescent microscope, and images captured using a camera and analyzed with ImageJ.

Identification of the transfected neurons on DAT^iCRE^ mice by immunohistofluorescence was performed as described above, with the addition of 1:500 Chicken-anti-GFP primary IgG (ab13970, Abcam) in the solution. A Goat-anti-chicken AlexaFluor 488 (1:500, Life Technologies) was then used as secondary IgG. Neurons labeled for TH in the VTA allowed to confirm their neurochemical phenotype, and those labeled for GFP to confirm the transfection success.

#### Model and Behavioral Potential Energy Theory overview

The details of the model can be found in the Supplementary Methods. In short, at its largest scale, the e-mouse model was designed as a distributed decision architecture deciding how an e-mouse navigates in a space. To fit experimental paradigms, we considered the physical space (a circular arena), but the model could extend to any task space. The e-mouse navigation was governed by linear speed and angular commands ensuring convergence toward either a default objective (circling along arena walls) or goal-directed behavior toward an internal goal, set by a recurrent prefrontal neural circuit. The contribution of default behavior to speed was high when the e-mouse headed toward, or was aligned with, the arena walls, but vanished when the e-mouse was far from, or not aligned with, the arena walls. Angular dynamics toward the default objective ensured that the e-mouse aligned with the wall when approaching it. Far from walls, angular dynamics were essentially influenced by goals situated in its visual foreground landscape. The internal goal was determined according to a probabilistic soft-max process (modeling basal ganglia operations), which stochastically selected the preferred position of neurons according to probabilities based on their instantaneous spiking rate. Neuronal preferred positions were organized on a square lattice that covered the arena.

The local recurrent prefrontal network consisted in a detailed biophysical model of PFC neurons and connections (*48*). The model contained neurons that were either excitatory (E) or inhibitory (I), with sparse connectivity, an E/I ratio of 4, and E/I current balance at the post-synaptic neuron level. Leaky integrate-and-fire (LIF) neurons were endowed with recurrent and feed-forward currents, and with adaptive action potential threshold in excitatory neurons. Feed-forward currents consisted of AMPA currents while recurrent currents consisted of AMPA, NMDA, GABA-A and GABA-B currents. We considered a uniform delay for synaptic conduction and transmission. AMPA feed-forward currents consisted in two parts: 1) inputs from external sources (putatively sub-cortical and/or cortical inputs), modeled as an exponentially-filtered normal stochastic process with temporally homogeneous mean_[MS1]_, and 2) hippocampal place-field inputs encoding the e-mouse position, with PFC neurons receiving input currents proportional to the activation of their receptive fields by a Gaussian input centered on e-mouse position. Recurrent NMDA currents were subject to dopamine modulation that affected their maximal conductance, in all synapses of the network (“DA-excitability”).

Network excitatory synapses underwent a dopamine (DA)-modulated form of Hebbian Spike Timing-Dependent Plasticity (STDP) (“DA-plasticity”), with pre- then post-synaptic spike sequences leading to long-term potentiation (LTP), and post-then pre-synaptic spike sequences to long-term depression (LTD). Spiking activity patterns did not translate into immediate effective synaptic changes, but rather resulted in synaptic tags, called eligibility traces (*20*), which were read out at the time of dopamine release (*49*). Eligibility traces (eLTP and eLTD, respectively) arose from synaptic calcium dynamics in the postsynaptic button (*48*, *50*). Synaptic calcium took into account the sum of calcium contributions arising from pre- and post-synaptic spiking, together with buffering and extrusion. Intracellular calcium activated calcium-dependent kinases and phosphatases, which competed to form eLTP and eLTD traces. Dopamine gated the transformation of eLTP and eLTD traces into actual changes in excitatory synaptic weights. Dopamine level was the same at all synapses. Dopamine was released when the e-mouse was detected inside rewarded areas, but also occurred spontaneously according to a Poisson process, i.e. with homogenous release probability within each time bin. The dopamine concentration followed second-order dynamics modeling release and recapture.

---

**Supplementary Figure 1:**
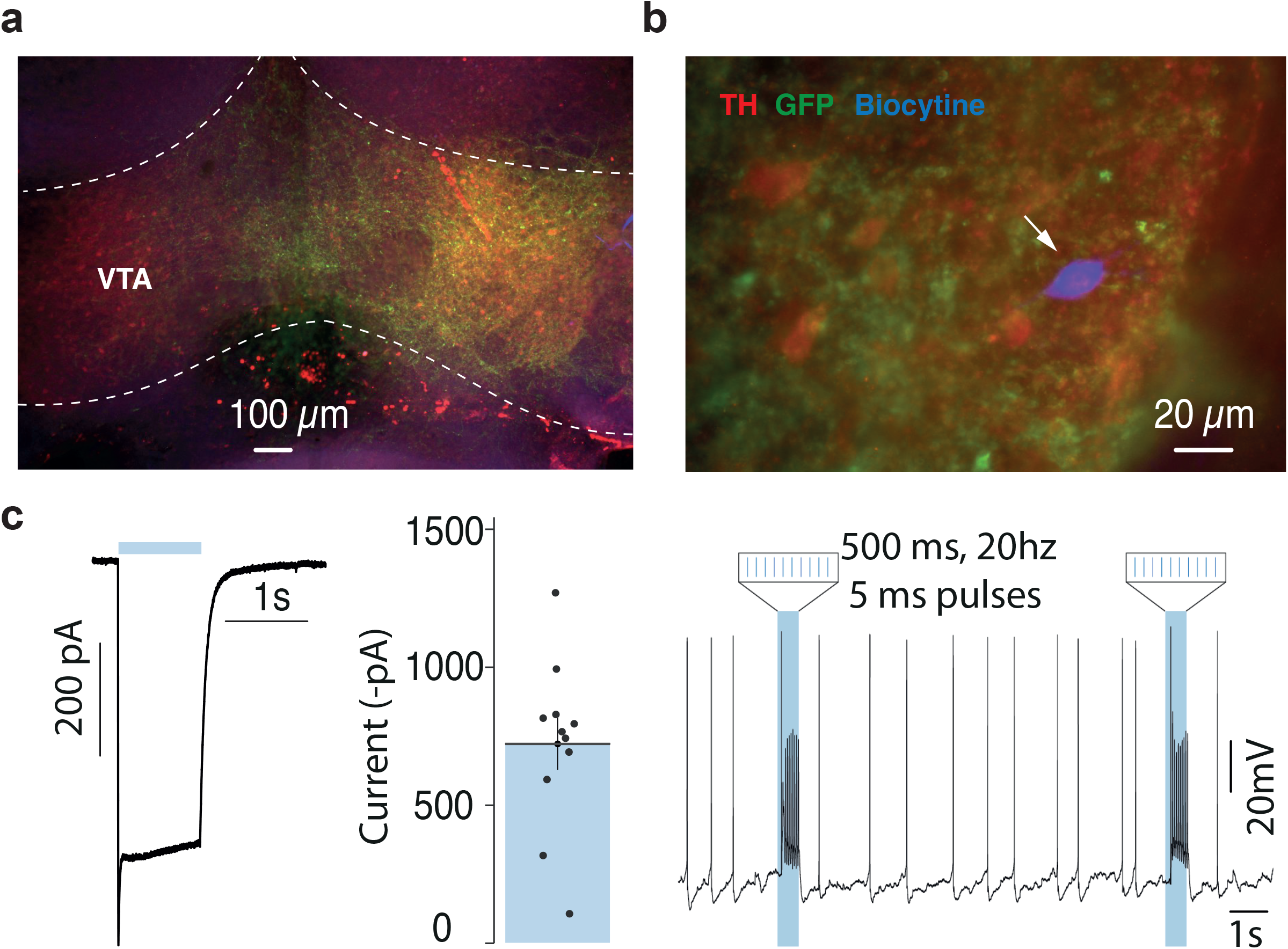
Specificity of dopamine control by optogenetics a. ChR2 was expressed in VTA DAT+ (dopamine) neurons in slices from DAT-Cre mice used for ex-vivo recording. b. Zoom in the example neuron recorded, expressing TH, YFP and filled with biocytin (blue). c. Left, example of current induced by a one second-pulse and average currents from 12 cells, induced by the 10 5ms-pulses at 20Hz. Right, example of bursting driven by 10 5ms-pulses at 20Hz

**Supplementary Figure 2:**
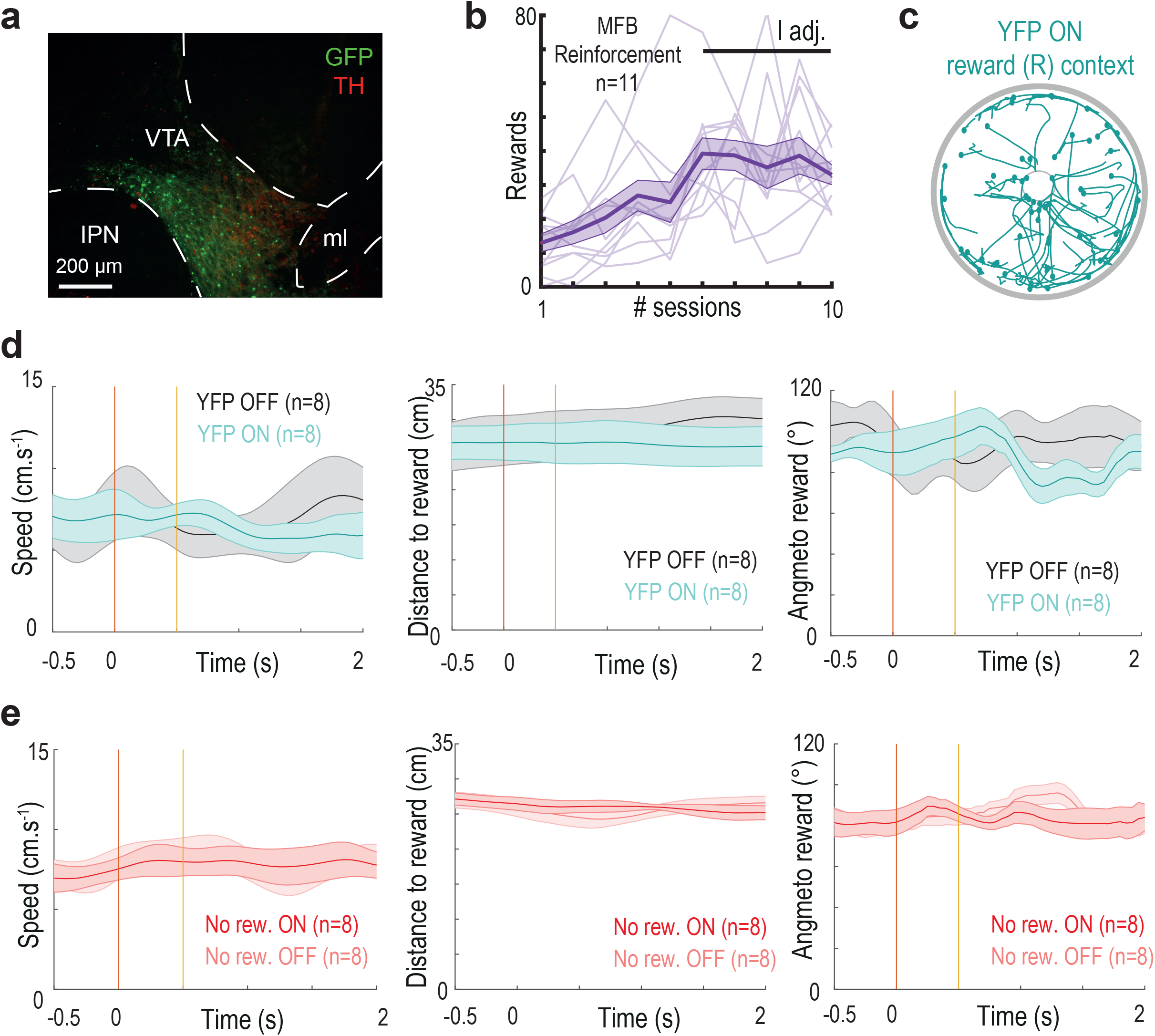
control experiments for Figure 3 a. ChR2 was expressed in VTA DAT+ (dopamine) neurons in animals used in Figure 3 experiments. b. Number of location visits across sessions of MFB reward learning. c. Post-photostimulation bouts of trajectories in the YFP, ON light, R context. d. From left to right: speed, distance and angle to rewarded location around the time of random VTA photostimulation in the periphery for YFP animals. e. Same as d for Chr2 animals in the “no reward” condition.

## Supplementary methods related to the MAGNet model, simulations and BPE theory

### e-mouse navigation

In the MAGNet model, e-mouse navigation was modeled, in a circular arena (radius *r_arena_*), as a process where orientation and speed were governed by a convergence toward either a default objective that consisted in approaching and aligning with the arena wall (answering to a need for security), or a goal-directed objective, answering to a need for exploration, the discovery and the retrieval of rewarded locations (i.e. circles with radius *r_reward_*). While the default behavior was set according to ballistic laws in the model, goals were driven by population dynamics of the recurrent neural network (see below).

The mouse position was denoted *P* = {*X_P_, Y_P_*}, with *X_P_* and *Y_P_* its cartesian coordinates. The position vector was

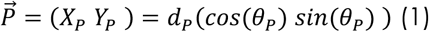

with 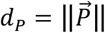 the distance to the center of the arena *0* and 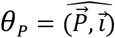 the directional angle of the position vector 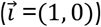. The mouse moved according to

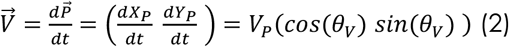

where *V_P_* was the linear speed and 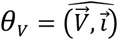 the direction of movement, i.e. the directional angle of the mouse speed vector, termed hereafter the speed angle.

### e-mouse linear speed dynamics

The e-mouse linear speed obeyed

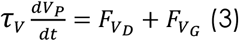

where the terms *F_V_D__* = *V_D_* – *V_P_* and *F_V_G__* = *V_G_* – *V_P_* modeled the contribution of default (subscript D) and goal behaviors (subscript G) to linear speed.

On the one hand, *F_V_D__* drove linear speed toward the default command speed *V_D_*, which was expressed as

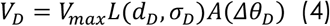

where *V_max_* was the maximal linear speed, 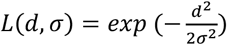 and 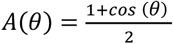 respectively denote exponential colinear (with characteristic distance *σ*) and cosine angular tuning functions for motor commands (*1*), *d_D_* the distance separating the e-mouse and the default objective *D*, and Δ*θ_D_* = *θ_V_* – *θ_D_* the angular difference between the speed and default objective angles.

At each time, *D* was defined as the nearest point from e-mouse’s position situated on a circle concentric with the circular arena wall with *r_d_* = *r_arena_* – *r_mouse_*, with *r_arena_* the arena radius and *r_mouse_* the e-mouse body’s half width, i.e. at the nearest possible distance from the wall, when considering the physical dimension of the e-mouse body. The default objective angle was computed as *θ_D_* = (1 – *L*(*d_D_, σ_D_*))*θ_P_* + *L*(*d_D_, σ_D_*)*θ_T_*, where *θ_P_* was the directional angle from the animal position *P* to its projection onto the wall *D*, and *θ_T_* was the directional angle tangential to the arena circular wall at point *D* and in the direction of e-mouse movement.

Overall, *F_V_D__* modeled the propensity of e-mouse to be driven by the default command speed *V_D_*, which was important when the e-mouse was 1) approaching the arena wall and heading toward it (typically small *d_D_* (yielding *θ_D_*~*θ_P_*) and *θ_V_~θ_P_*, resulting in substantial *L*(*d_D_, σ_D_*) and *A*(*Δθ_D_*) values) and 2) aligning parallel to the arena wall (typically *d_D_*~0 (yielding *θ_D_~θ_T_*) and *θ_V_~θ_T_*, resulting in large *L*(*d_D_, σ_D_*) and *A*(*Δθ_D_*) values). Conversely, the contribution of the default behavior to the e-mouse overall speed vanished when the e-mouse was far from, or not aligned with, the arena wall.

On the other hand, *F_V_G__*, drove the e-mouse linear speed toward the goal command speed *V_G_*, which was expressed as

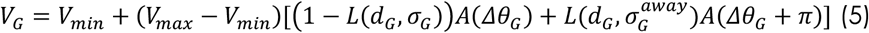

where *V_min_* was the e-mouse’s minimal linear speed, *d_G_* the distance separating the e-mouse and goal objective (hereafter denoted as the internal goal) *G*, *Δθ_G_* = *θ_V_* – *θ_G_* the angular difference between the speed angle and 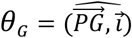 the directional angle from the e-mouse to the internal goal. Altogether, *F_V_G__* modeled the propensity of the e-mouse to be driven by the goal command speed, which was important when the e-mouse was 1) far from the internal goal and heading toward it (large *d_G_* so (1 – *L*(*d_G_, σ_G_*)~1 and *θ_V_ ~θ_G_* such that *A*(*Δθ_G_*) is large), or 2) nearby the internal goal and moving away from it (small *d_G_* so 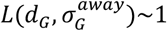 and *θ_V_ ~θ_G_* + *π* such that *A*(*Δθ_G_* + *π*) is large). The scaling of linear tuning functions, when moving toward or away from the internal goal were determined by *σ_G_* and 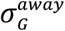, with 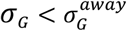 so that navigation was faster when escaping away from a recently visited rewarded point. This hypothesis was necessary to avoid otherwise inevitable (although unrealistic) e-mouse repeated navigational loops at rewarded locations.

The internal goal *G* was determined according to a probabilistic soft-max process with *G* drawn, at each time-step, from the normalized exponential probability distribution

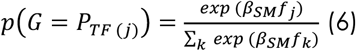

where 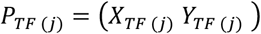 was the preferred position and *f_i_* the estimated firing frequency of neuron *j, β_SM_* the inverse temperature of the process and *k* indexing neurons taking part to the soft-max. The estimated firing frequency was obtained by filtering spiking with an exponential kernel with time constant *τ_F_*. Preferred positions were organized on a square lattice following the *X* and *Y* axes that covered the arena, with 70% of neurons within the arena and 30% outside on *X* and *Y* axes. Neurons’ preferred positions covered a surface area more than twice that of the arena, so that the internal goal could lay outside the arena. The soft-max was thus computed with neurons whose preferred positions were closer than 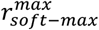, so that *G* essentially laid within the arena. This ensured that goal-directed influence balanced the centrifugal influence of the default behavior, such that a naïve (i.e. before learning) e-mouse spent ~60% of their time in the default behavior (i.e. running along walls). Convergence to the internal goal could nevertheless tend to drive the e-mouse outside the arena sometimes. To avoid this unrealistic behavior, the distance of the e-mouse to the arena center, *d_P_*, was reset to *r_d_* when this happened.

### e-mouse angular dynamics

The e-mouse angular direction *θ_V_* obeyed

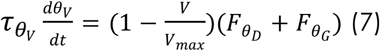

with the first term catching the slower rotation of animals when moving faster, and *F_θ_D__* and *F_θ_G__* represented contributions of default and goal behaviors to e-mouse orientation changes.

Rotation speed toward the default objective was governed by

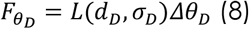

so that it was larger when *θ_V_* was far from *θ_D_* and when the e-mouse approached arena walls (*L*(*d_D_*, *σ_D_*)~1), which insured a progressive rotation toward *θ_T_* (i.e. the e-mouse aligned with the wall when approaching). Rotation was essentially independent of the default behavior far from the wall, being instead mostly goal-directed, with rotation toward the internal goal obeying

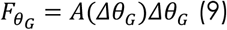

where rotational speed scaled with the difference between e-mouse’s direction *θ_V_* and *θ_G_* the angle facing the internal goal, but only when the e-mouse was essentially influenced by goals situated in its visual foreground landscape (*A*(*Δθ_G_*) vanished at large *Δθ_G_* values). This hypothesis, which expressed a visual gating of internally-guided behaviors, reduced the noise of goal-directed navigation but was not essential to the results.

### Pause and redirection behaviors

The e-mouse had behavioral pauses (during which rotational or linear speed was null) that occurred spontaneously with increasing probability when closer to the arena wall, as in real mice. Pause times where thus drawn according to a Poisson process with a rate scaled with the distance to the center of the arena: 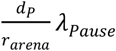, each pause lasting *d_Pause_*. Redirections of the e-mouse occurred at the end of pauses, by drawing the new angular direction from a von Mises distribution (*2*) with mean *θ_V_* and concentration *κ_redir_* (i.e. with a circular standard deviation of 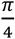). In order to avoid unrealistic redirections toward the exterior of the arena when at its edges, directions were redrawn when 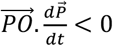 (centrifugal redirection) with probability *p_redraw_* = *L*(*d_D_*, *σ_D_*) (nearby 1 in the close vicinity of the arena wall).

### Local recurrent neural network biophysical model

We built a biophysical model of a prefrontal local recurrent neural network, endowed with detailed biological properties of its neurons and connections (*3*). The network model contained *N* neurons that were either excitatory (E) or inhibitory (I) (neurons projecting only glutamate or GABA, respectively (*4*)), with probabilities *p_E_* and *p_I_* = 1 – *p_E_* respectively and 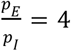 (*5*). Connectivity was sparse (i.e. with probability connection *p_c_* (*6*)), with no autapse (self-connections). Synaptic weights *w*_(*i,j*)_ of existing connections were initiated with a value *μ_w_*, before possible consecutive additional Hebbian assemblies were learnt or written by hand (see below).

To cope with simulation times required for the massive explorations of the model, neurons were modeled as leaky integrate-and-fire (LIF) neurons. The membrane potential of neuron *j* obeyed

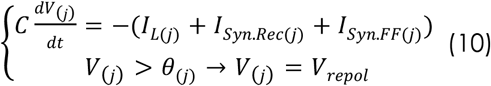

where was *V_rest_* the repolarization potential. The action potential (AP) threshold *θ_(j)_* was adaptive in excitatory neurons, with spike-induced instantaneous increase and exponential convergence with time constant *τ_θ_* toward its steady-state value *θ*_0_:

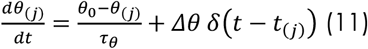

where *δ* represents the Dirac function and *t*_(*j*)_ AP times in neuron *j*.

The leak current followed

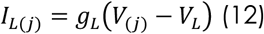

with *g_L_* the leak conductance and *V_L_* its equilibrium potential.

The recurrent synaptic current on postsynaptic neuron *j*, from either excitatory or inhibitory presynaptic neurons (indexed by *i*), was

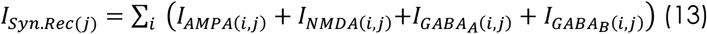

The delay for synaptic conduction and transmission, *Δt_syn_*, was considered uniform across the network (*7*). Synaptic recurrent currents followed

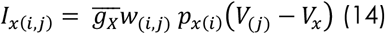

Where 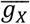 was the maximal conductance, *w*_(*i,j*)_ was the synaptic weight, *p*_*x*(*i*)_ the opening probability of channel-receptors and *V_x_* the reversal potential. The NMDA current followed specific dynamics

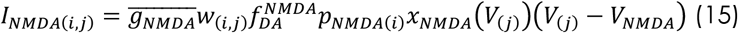

accounting for the voltage-dependence of the magnesium block (*8*) which was modeled as

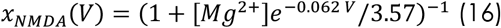

and 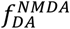 represented the dopamine-dependent gating of NMDA conductance (*9–11*) through D1-receptors, affecting equally all synapses of the network (diffuse VTA dopamine input), according to

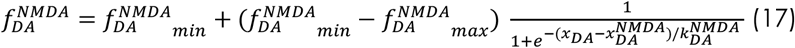

where 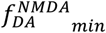 and 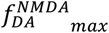 set minimum and maximal gating and 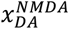 and 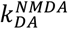 were the half-activation and inverse slope of DA concentration sigmoidal effect.

AMPA and GABAA channel rise times were approximated as instantaneous (*7*) and bounded, with first-order decay

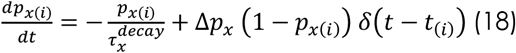

where *t*_(*i*)_ represented the pre-synaptic APs’ times. In order to account for the longer NMDA (*12*) and GABAB (*13*) channel rise times, opening probabilities followed second-order dynamics (*7*)

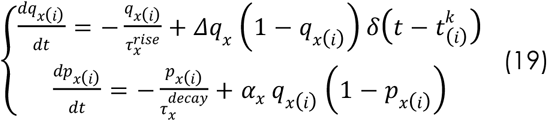

Recurrent excitatory and inhibitory currents were balanced on average in post-synaptic neurons (*14*) according to driving forces and excitation/inhibition weight ratio, through

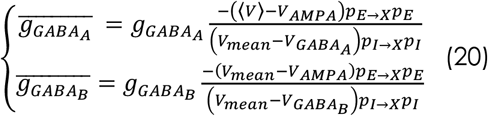

with 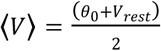 an approximation of the average membrane potential, and *X* the excitatory or inhibitory identity of the postsynaptic neuron receiving the inhibitory current.

The feed-forward synaptic current *I*_*Syn.FF*(*j*)_ – putatively arising from sub-cortical and/or cortical inputs – consisted of an AMPA current

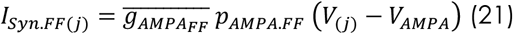

where *p_AMPA.FF_* was the sum of two components,

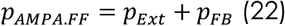

The first one, *p_Ext_*, corresponded to network-wide AMPA inputs from external sources for every network neuron, and built as the convolution by an exponential kernel *k_Ext_* (time constant *τ_AMPA_*) of a random stochastic process drawn from the normal distribution, with mean and standard deviation derived from the binomial distribution of the number of input spikes per time step when considering *n_Ext_* external independent inputs projecting onto the network and a spiking probability *x_Ext_* for each input per given time step:

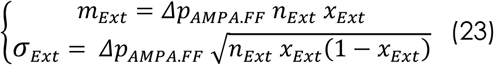

The second component, *p_FB_*, corresponded to the putatively hippocampal feedback encoding the e-mouse position (Figure 1e), with neuron *j* receiving an input current proportional to the activation *k_FB_*(*j*) of their preferred position. Activation function were modeled as bivariate distributions centered on the preferred position (*X*_*RF*(*j*)_, *Y*_*RF*(*j*)_).

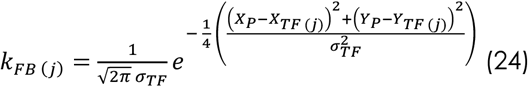

which displayed similar, but flatter profiles, compared to bivariate distributions, to insure a more homogeneous feed-back activation of neurons encoding the e-mouse position and, as a consequence, smoother and more stable learning (see below). Activation function width was determined by *σ_TF_*. Finally, the feed-back opening probability was *p_FB_* = *k_FB_ x_FB_*, with *x_FB_* a constant.

### Synaptic plasticity

We built a constrained biochemical model of the pathways’ architecture implicated in the dopaminergic reinforcement of synaptic plasticity. Network excitatory synapses underwent a dopamine (DA)-modulated form of Hebbian Spike Timing-Dependent Plasticity (STDP), with pre-then post-synaptic spike sequences leading to potentiation (and post-then pre-synaptic spike sequences depression). Spiking activity patterns did not translate into immediate effective synaptic changes, but rather resulted in synaptic tags, called eligibility traces (*15*), which were read out at the time of dopamine release (*16*). Standard Hebbian synaptic STDP rules devoid of reinforcement gating would strengthen any e-mouse navigation trajectory associated with a chain of neuronal activation. By contrast, in the presence of DA-reinforced plasticity, network synapses are only modified if they participated to rewarded trajectories.

At the molecular scale, the spike timing-dependence of synaptic plasticity (*17*, *18*) was considered to arise from synaptic calcium dynamics in the postsynaptic button (*3*, *19*). Specifically, calcium was computed as

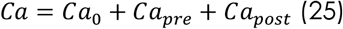

which took into account calcium the sum of calcium contributions arising from pre- and postsynaptic spiking. Presynaptic calcium dynamics followed

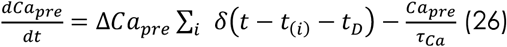

which modeled the calcium influx due to pre-synaptic spiking through Voltage-Dependent Calcium Channels (VDCCs), with *ΔCa_pre_* the calcium step per action potential (AP), *t*_(*i*)_ APs’ times, and *t_D_* the delay necessary for AMPA channels’ activation and excitatory postsynaptic potential (EPSP) buildup driving VDCCs’ opening), in addition to extrusion/buffering of this calcium source, with time constant *τ_Ca_*. Postsynaptic calcium dynamics followed

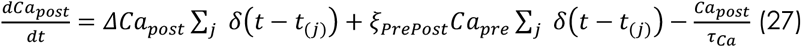

which took into account extrusion/buffering (last term) in addition to the calcium influx from post-synaptic back-propagated spiking opening VDCCs (first term) and NMDA channels (second term). The NMDA calcium influx was scaled by an interaction coefficient *ξ_PrePost_* and depended on the product of the presynaptic calcium contribution and postsynaptic spiking, to account for the associative opening of NMDA channels due to magnesium blockade.

Intracellular calcium activated calcium-dependent kinases and phosphatases (putatively, CaMKII kinase and calcineurin) that competed to form molecular traces (*18*, *20*, *21*), i.e. eligibility traces (*16*, *21*, *22*), and which were distinct for potentiation (eLTP) and depression (eLTD) processes (*18*). These traces putatively competed for the phosphorylation of an ERK tag (*22*–*24*), which would decay to a non-phosphorylated state if not consolidated by dopamine into effective – reinforced – changes in synaptic weights. In the model, each eligibility trace followed first-order dynamics, i.e.

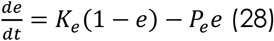

where kinase and phosphatase activation followed

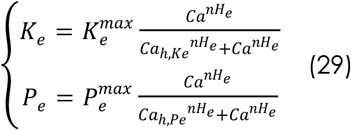

with 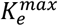 and 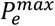 the maximum rates, *Ca_h,Ke_* and *Ca_h,Pe_* the half-activation calcium values and *nH_e_* the Hill coefficient

Experimental studies indicate that the activation of D1 receptors by dopamine increases cAMP levels and, consequently, protein kinase A (PKA) activity (*11*, *21*, *23*), resulting in the transformation of eligibility traces into effective – reinforced – synaptic changes (*16*, *25*), i.e. modified glutamate receptor densities or phosphorylation levels, e.g. through CREB-induced protein synthesis (*23*, *24*). In the model, excitatory synaptic weights *w* evolved according to a dopaminergic gating of a kinase/phosphatase cycle activated by *e_LTP_* and *e_LTD_* eligibility traces (*18*), with first-order (soft-bound) kinetics :

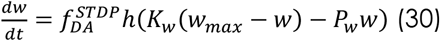

with *w_max_* corresponded to the maximal synaptic weight, 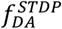 the dopamine-gated functional fraction of the kinase/phosphatase cycle and *h* a variable accounting for homeostatic synaptic regulation required only for online learning simulations (see below). Kinase and phosphatase activations followed

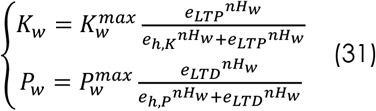

with 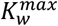 and 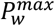 the maximum rates, *e_h,K_* and *e_h,P_* the half-activation eligibility values and *nH_w_* the Hill coefficient.

The dopaminergic gating of synaptic plasticity operated on all synapses (diffuse VTA dopamine input) through D1-receptors (*10*, *11*), and followed

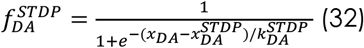

where 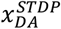 and 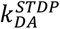 were the half-activation and inverse slope of DA concentration sigmoidal effect on plasticity.

### Dopamine dynamics

The dopamine concentration, following spontaneous or reward events (at time *t_DA_*), obeyed second-order dynamics

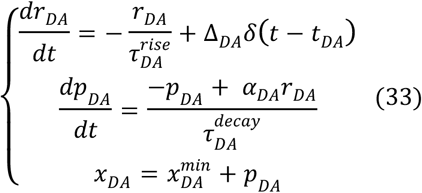

where 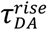 and 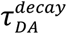 were rise and decay time constants, 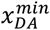 the minimum DA concentration, and *α_DA_* a parameter scaling the influence of *r_DA_* on *p_DA_* dynamics and adjusted to get a maximal value 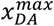. Spontaneous events were drawn according to a Poisson process with a rate 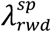, with a refractory period 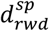. Reward events occurred when the e-mouse entered a rewarded location. In simulations with three rewarded locations, following consecutive visits of the same location were not rewarded.

### Numerical procedures and parameters

The MAGNet model was simulated and explored using custom developed MATLAB code, whose differential equations were numerically integrated using the forward Euler method (*Δt* = 1*ms*). Most simulations, achieved in offline conditions (Figure 2), were achieved with the following set of standard parameter values: space and navigation: *τ_V_* = 500 *ms*, *τ_θ_V__* = 50 *ms*, 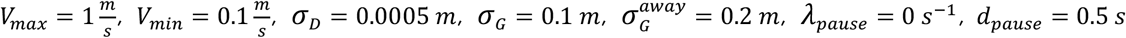, 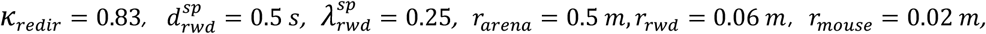 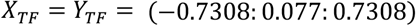; neural decoding into the internal goal : *β_SM_* = 1.5, *τ_F_* = 100*ms*; neural encoding of the e-mouse position : *σ_RF_* = 0.075 *m*, *x_FB_* = 0.075; network architecture: *N* = 500, *p_E_* = 0.8, *p_c_* = 0.75, *Δt_syn_* = 1*ms*, *μ_w_* = 0.1, *σ_w_* = 0, *w_max_* = 5; intrinsic properties, *C* = 1 *μF. cm*^−2^, 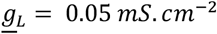, *V_L_* = −70 *mV*, *θ*_0_ =−50 *mV*, Δ*θ* = 50 *mV*, *τ_θ_* =50 *ms*, *V_repol_* =−60 *mV*; recurrent currents: 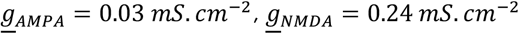, *g_GABA_A__* = 0.03 *mS. cm*^−2^, *g_GABA_B__* = 0.00003 *mS. cm*^−2^, *V_AMPA_* = *V_NMDA_* = 0 *mV*, *V_GABA_B__* = −70 *mV*, *V_GABA_B__* = −90 *mV*, [*Mg*^2+^] = 1.5 *mM*, 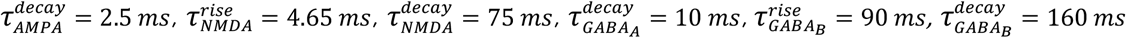, *α_NMDA_* = 0.275 *ms*^−1^, *α_GABA_B__* = 0.015 *ms*^−1^. *Δp_AMPA_* = *Δq_NMDA_* = *Δp_GABA_B__* = *Δq_GABA_B__* = 0.1; feed-forward currents : *p_Ext_* = 0, 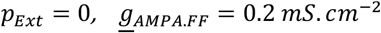, *f_Ext_* = 25 *Hz*, *n_Ext_*, = 30, *Δp_AMPA.FF_* = 0.1; calcium dynamics: *Ca*_0_ = 0.1 *μM*, *τ_ca_* = 50 *ms*, *ΔCa_pre_* = Δ*Ca_post_* = 0.5 *μM*, *ζ_PrePost_* = 6.5, *t_D_* = 20 *ms*; synaptic weight plasticity: 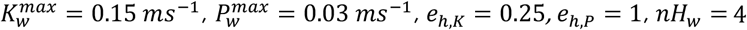; eligibility traces : 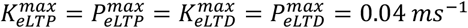, *Ca_h,KeLTP_* = 1.65*μM*, *Ca_h,PeLTP_* = 0.495*μM*, *Ca_h,KeLTD_* = 1.25 *μM*, *Ca_h,PeLTD_* = 0.375 *μM*, *nH_e_* = 4; dopamine properties: 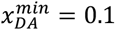, 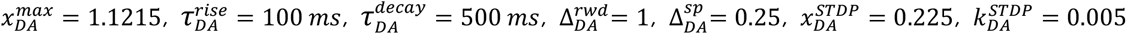, 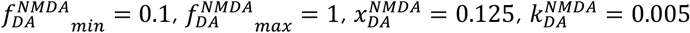.

### Initial conditions and simulation setups

The model was initialized with randomized membrane potentials (uniformly distributed in [*θ*_0_, *θ*_0_ − 5]*mV*) and synaptic channel openings mimicking average channel openings (*p_AMPA_*~0.0025, *p_NMDA_*~0.2, *p_GABA_B__*~0.0025, *p_GABA_B__*~0.15), as well as with e-mouse at initial random positions at distance *d_P_* = 0.75*r_d_*, null linear speed and random initial direction *θ_V_*.

Online learning simulations (Figure 1f) lasted 300 seconds and consisted of 10 concatenated simulations of 30 seconds termed sessions, with behavioral pauses when rewarded, without redirection. Online learning dynamics easily yielded saturated synaptic weights and neural activity, even with slower learning kinetic parameters 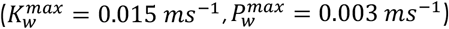. Such plasticity/activity runaway is a classical issue when assessing online learning in random recurrent networks. It arises from the positive feedback linking excitatory activity and plasticity between excitatory neurons and is likely stabilized by different homeostatic processes providing counteracting negative feedbacks at the neuronal and network scales. In the present decision architecture, this problem was largely amplified by the additional positive feedbacks linking connectivity and neural activity, on the one hand, and emice behavior, on the other hand. For instance, increased connectivity at rewarded locations (Hebbian assemblies) increased reward rates, which in turn increased DA-reinforcement of STDP at these Hebbian assemblies. In the context of the present study, we found that synaptic homeostasis was essential in the online learning setup (Figure 1f). Therefore, synaptic homeostasis was constrained by distinct homeostatic processes at excitatory synapses, which were required in online simulations, but not offline simulations (Figure 1g and 2; see below). Hence, in addition to a hard-bound *w_max_* = 5, excitatory-excitatory synapses underwent synaptic scaling, which normalizes synaptic connections. We considered a form of synaptic scaling that included both presynaptic and postsynaptic normalization, i.e.,

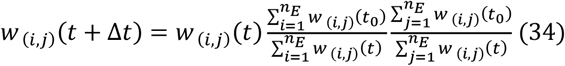

which allowed a limit to catastrophic plasticity/activity runaway eventually occurring at synapses linking neurons of Hebbian assemblies and the rest of the network. In addition, we considered two forms of saturation constraining plasticity runaway. First, by assuming that no plasticity occurred at spiking post-synaptic frequencies superior to a critical frequency set as *f_h_* = 1/*D*, i.e. putatively through presynaptic calcium saturation. This process helped avoiding plasticity/activity runaway within each Hebbian assembly. Second, by assuming that the total amount of post-synaptic potentiation admits a maximum within each neuron, putatively due to upstream resources availability (e.g. pool of precursors limiting the synthesis of new glutamatergic receptors). This process constrained the spatial extension of Hebbian assemblies. Altogether, these saturations terms wrote

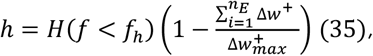

with *H* the Heaviside function, 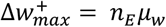 the maximal amount of possible potentiation changes and 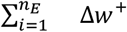 hard-bound limited by 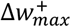. Although not crucial for offline learning (Figure 1g and 2), these homeostatic processes were kept in that case, for the sake of simplicity. In a similar vein, because getting strong and compact Hebbian assemblies during online learning proved difficult under certain modeling and parameter choices, we found easier to set constant eligibility time constants (which otherwise non-linearly depended on the calcium activation of their kinase/phosphatase cycles *τ_e_* = 1/(*K_e_* + *P_e_*)), with *τ_eLTP_* = *τ_eLTD_* = 250*ms*. For the sake of simplicity, this option was also kept for offline learning, although not essential in that setup (Figure 1g and 2).

Offline simulations (Figure 1g) mean performance rate were computed over 10 simulations in each of 18×18 conditions DA-excitability and DA-plasticity. Simulations lasted 60 seconds, with a pause rate *λ_Pause_* = 1/3 *s*^−1^. DA-excitability was parameterized with 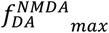 in the interval = [0,1]. DA-plasticity was mimicked by initiating the connectivity matrix before model simulation, as if plasticity had previously built three Hebbian assemblies (i.e. there was no plasticity during offline simulations). This initialization consisted in adding three Hebbian assemblies centered at rewarded locations. Each of these bivariate gaussian Hebbian assemblies consisted of the synaptic matrix *w_k_*, built as the auto-association (i.e. external product) of a vertical vector specifying gaussian distance of each neuron preferred position to rewarded location *k* (with spatial standard deviation *σ_rwd_* = 0.125 *m*), i.e.

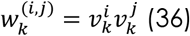

with

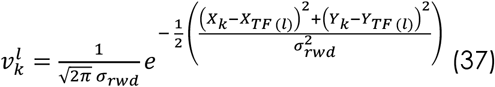

and *P_k_* = (*X_k_ Y_k_*) is the position of reward location *k*. In Figure 1f, we only kept synapses oriented toward each reward locations within Hebbian assemblies (i.e. *w*_(*i,j*)_ = 0 when *w*_(*j,i*)_ > *w*_(*i,j*)_) to assess a scheme where STDP would have yielded purely asymmetric connections, but similar results could be obtained with symmetric connections (not shown). The Hebbian assemblies were then normalized to have a maximal weight *w_max_* in the interval [0,5], and added to a constant mean *μ_w_*

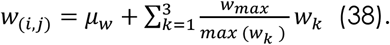

Offline learning simulations (Figure 2b) consisted of 100 successive learning trials lasting 1.5 seconds. To speed up simulations, e-mice were initialized with *d_P_* = 0.3 *m*, heading toward the central rewarded location and with maximum linear speed. In Figure 2d-f and 2h, simulations lasted 2.5 seconds, with the e-mouse initialized randomly in the arena, with maximum linear speed and random initial direction *θ_V_*. In Figure 2f, the angular speed was slower by a factor 10 before phasic DA, so that the e-mouse displayed equivalent positions at that time.

Realizations of von Mises distributions were numerically computed using the code developed by D. Muir (26).

### Behavioral Potential Energy (BPE) theory

In order to better understand how e-mouse behavior arises from past dopaminergic reinforcement (DA-plasticity) and online motivational dopaminergic modulation (DA-excitability), we built a simplified theory capturing essential causal and dynamical traits governing the full decision architecture model. To reduce dimensionality, we consider that, thanks to revolution symmetry in the one rewarded location setup, spatial behavior is reducible to one dimension, with rewarded location set at position *p_AH_* = 0. Also, the theory is built as a simplified representation of e-mouse navigation that neglects the detailed dynamics of linear and angular speed ballistic commands considered in the model. In particular, the contribution of the default behavior to linear speed, which is negligible at a certain distance from the arena walls in the model, is not taken into account. Moreover, we focus on how e-mouse navigation depends on the essential interactions linking *p*, the e-mouse position encoded by feed-forward hippocampal inputs to the network resulting in a bump of neural activity (see e.g. Figure 1e, lower panel, top maps), and *g*, the internal goal position decoded downstream by basal ganglia through soft-max computation.

This theoretical framework illustrates how goal-directed mouse behavior can be interpreted in the framework of attractorial dynamics within a landscape of behavioral potential energy (BPE), which depends on both DA-plasticity and DA-excitability. Building the theory unraveled two mechanisms driving e-mouse navigation. The first mechanism relates to the local positional stability of the activity bump, in the vicinity of the Hebbian assembly (HA). The second mechanism acts at a global scale of the arena and depends on the neural activity at the HA. Thus, in the theory, we posit that e-mouse *p* and DA dependent goal-directed navigation obeys a velocity law including the two mechanisms

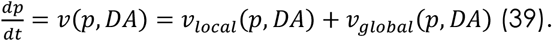

In the following, we first assess how each term can be described in a reduced and tractable fashion from the model dynamics. We then show how BPE can be derived and interpreted.

The first, local, mechanism relates to the positional stability of the activity bump. In the case when no HA is present (i.e., before reward-place learning; e.g., Figure 2b, trial #1, left panel), both feed-forward inputs encoding the e-mouse position and recurrent connections are symmetric with regard to position *p*, such that the activity bump displays a symmetric spatial firing frequency around *p*. As a result, *g*, the decoded internal goal position, which statistically reflects the position of the activity bump maximum, is also situated at *p*. Hence, the e-mouse is on its goal, convergence is already achieved and there is no movement.

By contrast, let’s consider the case where a HA is present (due to previous place-reward association; Figure 2b, trial #100, left panel), with the e-mouse in the vicinity of the HA. In such a situation, within the activity bump, excitatory synaptic currents generated in excitatory neurons closer to the HA, by excitatory neurons farther from the HA (centripetal currents), are larger than centrifugal currents generated at reciprocal synapses. This is due to the fact that centripetal currents occur at synapses with larger synaptic weights (i.e., higher in the HA weight gradient), compared to centrifugal currents. The resulting firing frequency profile of the activity bump is biased toward the HA, so that, on average, the decoded internal goal position, *g*, lies closer to the HA, compared to *p*. As *p* converges toward *g*, it continuously moves in the direction of the HA. In turn, as *p* is moving toward the HA, so too do the activity bump and its soft-max readout, *g*. Altogether, the weight gradient yields an attractorial convergence of the activity bump, *g* and *p* toward the HA. This convergence will obviously be stronger near the HA, where the synaptic gradient is steeper. Moreover, the large increase in NMDA currents mediated by DA (DA-excitability) will strongly amplify the gradient of excitatory currents due to DA-plasticity (i.e., the difference between centripetal and centrifugal currents) and the subsequent attractorial convergence toward the HA, through a *deepening* of the HA attractor, as unraveled by the theory (see below and Figure 2g-i). Accordingly, the local mechanism to e-mouse velocity scales with the gradient of excitatory currents

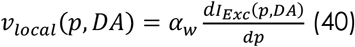

where *α_w_* is a constant and *I_Exc_*(*p,DA*) is the DA-dependent excitatory current received by excitatory neurons at position *p*, which can be approximated as

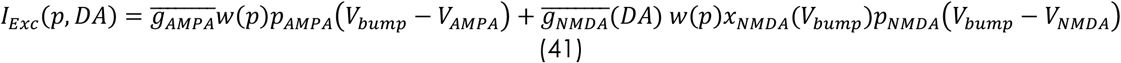

where 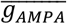 and 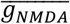 are maximal conductances with 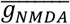 depending on DA-excitability, *w*(*p*) the sum of incoming synaptic weights on the neuron with preferred position centered at position *p*, *x_NMDA_*(*V_bump_*) is the non-linear activation of NMDA channels at the mean bump membrane potential *V_bump_* and where *p_AMPA_* and *p_NMDA_* gating variables at firing frequency *f_bump_* can be obtained by steady state approximation from equations 18 and 19 :

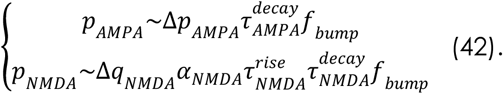

Therefore,

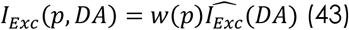

with

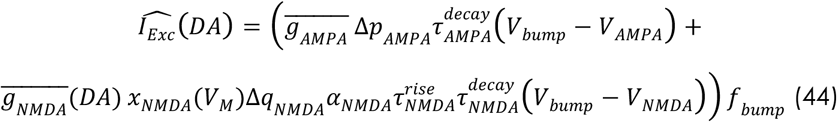

is the DA-dependent current per weight unit at firing frequency *f_bump_*. Note that 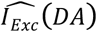 is an inward current, i.e., algebraically negative, which yields the sign of the local contribution to velocity and Behavioral Potential Energy (equations 40 and 56, see below). Note also that inhibitory currents can be neglected, as they display no spatial weight gradient in the model. The local mechanism contribution thus writes

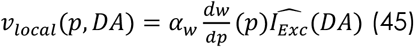

and depends on both DA-plasticity (i.e., on the weight gradient) and DA-excitability (i.e., DA-modulated NMDA current in the activity bump).

The second mechanism acts at the global spatial scale and also emerges from the interaction of DA-plasticity and DA-excitability: it arises from the DA-dependent increase of NMDA currents within the HA itself. Generally, the internal goal *g* is detected at *p* because the activity bump is the strongest spot of activity in the network. However, in the presence of DA, the increase of NMDA currents is boosted by large HA weights, which triggers massive associative co-activation of neuronal activity in the HA. Therefore, *g* almost instantaneously switches to 0, the AH position (see Figure 2f center panel and Figure 2h center column). Note that, due to noise in network dynamics within model simulations, activity can still be higher at *p* than at *g* in a number cases, accounting for why *g* does not always converge to the HA (Figure 2h center column). When acting, this mechanism operates at the global scale of the whole arena, independent of the position of the e-mouse (by contrast to the first mechanism, which acts locally in the vicinity of the HA). Thus, it yields attractorial convergence of the activity bump toward the HA through a widening of the HA attractor (as opposed to attractor deepening in the local mechanism), as shown below (see Figure 2g-i).

So, the global velocity contribution writes

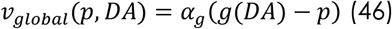

Here, for the sake of simplicity, we use a crude linear dependance of the distance of the emouse to the internal goal. However, using more complex dependance reminding model ballistics – or even zero order dependance – would yield qualitatively similar results. The essential point here is that, as shall be seen below, BPE increases with distance in all these cases. Moreover, based on simulations, we set *g*(*p, DA* = 0)~*p* when DA is absent. By contrast, when DA is present, decoded internal goal position *g*, is statistically essentially detected at either one of the two higher spots of activity in the network, i.e., the activity bump (*p*) and the HA (*p_AH_*), with probabilities related to their respective spiking frequency. Approximately, the internal goal position *g* can thus be estimated to lie, on average, at the barycenter of both spots weighted by their spiking activity

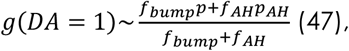

when DA is present. In that case, the global velocity therefore writes

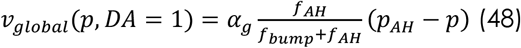

Setting

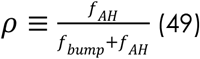

and leveraging on the fact that *p_AH_* = 0 leads to

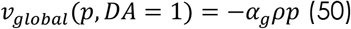

when *DA* = 1. Lumping both cases (*DA* = 0 and *DA* = 1) is possible by writing:

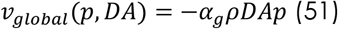

Overall, the velocity law governing e-mouse *p* and DA dependent goal-directed navigation is thus

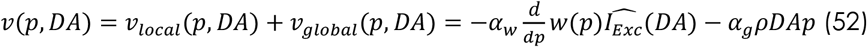

In the case where both DA plasticity and DA-excitability are present and the e-mouse is in the vicinity of the HA, the local and global terms hypothesize distinct positions of *g*, i.e., at *p* or *p_AH_*, respectively. However, in that case, *p_AH_* and *p* are practically almost confounded in the context of the noisy chaotic activity of the network. Moreover, and as a consequence, the global effect is minute, compared to the local effect, which is overwhelming. We therefore kept this crude formulation for the sake of simplicity, without developing a more complex description taking into account which term has to be considered in which case (presence or not of DA-plasticity and DA-excitability, distance to the HA).

The potential of any one dimensional dynamical system

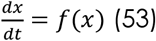

can be computed as

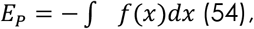

based on the physical idea of the potential energy (*27*). We therefore define the Behavioral Potential Energy (BPE) of the e-mouse at each point as

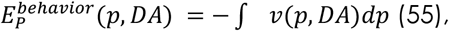

which yields

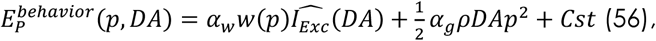

where *Cst* is an integration constant.

This expression captures how, at a previously rewarded location, reinforced HA weights induce attractorial dynamics though both their profile, whose gradient locally destabilizes the activity bump, and their strength, which shifts the internal goal at the global scale. Moreover, this expression accurately accounts for how shape, width and depth of the Hebbian-based attractor depend on previous DA reinforcement (DA-plasticity), current DA motivation (DA-excitability) and their interaction, and how it acts depending on e-mouse position. In doing so, it offers a framework for interpreting coupled dynamics between collective network activity at the activity bump and the HA, the e-mouse position, and the internal goal. Specifically, it mechanistically accounts for weak convergence to the – latent – attractor at the previously rewarded location under DA-plasticity alone (Figure 2f-i, left), deepening and widening of attractorial convergence under DA-plasticity + DA-excitability (Figure 2f-i, center and Figure 2h) and the absence of attractorial convergence under DA-excitability (Figure 2f-i, right).

BPE was computed using a gaussian-shaped distribution of weights

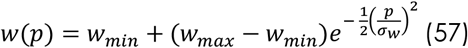

centered at position 0 and with spatial standard deviation *σ_w_*. For purely illustrative purpose in Figure 2, a phenomenological term was added to BPE to account for short-distance attraction to arena walls due to the default behavior:

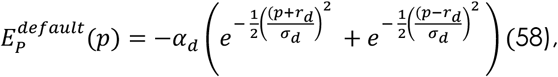

but this term is not part of the theory by itself. Regarding display specificities, in Figure 2i, onedimensional BPE was integrated in the range [*r_−arena_, r_arena_*], using integration constants chosen so that 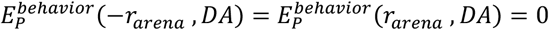 in each condition (DA-plasticity, DA-excitability, DA-plasticity + DA-excitability). In Figure 2h, BPE (left column) was generated in two dimensions from one-dimensional BPE (Figure 2i) by revolution symmetry, for the sake of illustration, i.e., visual correspondence with model simulations (center and right columns). BPE contour levels correspond to *BPE* = −[0 0.05 0.1 0.2 0.4 0.6 0.8 1]. Theory parameters were as following: *w_min_* = 0.1, *w_max_* = *w_min_* in the DA-excitability conditions and *w_max_* = 3 in DA-plasticity and DA-plasticity + DA-excitability conditions. The firing frequency *f_bump_* = 16 Hz was derived from simulations. The mean voltage at the peak of the activity bump was also taken from simulations: *V_bump_* = [−32.5, −32.5, −25]mV in all conditions (such depolarized values in the bump arise from spiking-induced depolarization of the adaptive AP threshold). The Gaussian widths were *σ_w_* = 0.075*m* and *σ_d_* = 0.01*m*. In Figure 2, we used *α_w_* = 20*C*^−1^*m*^2^, *α_g_* = 1.25*s*^−1^,*α_d_* = 0.025*m*^2^*s*^−1^, but these parameters can be arbitrarily scaled without any qualitative change in the BPE landscape. We made no specific hypothesis concerning the relative values of firing frequency and considered the parsimonious case where *f_bump_* = *f_AH_*, i.e., *ρ* = 1/2 in Figure 2. Again, this specific choice had no consequence on the BPE landscape. Other theory parameters were as in model simulations.

## REFERENCES

1. W. Schultz, Behavioral dopamine signals. Trends Neurosci. 30, 203–210 (2007).

2. A. Westbrook, T. S. Braver, Dopamine Does Double Duty in Motivating Cognitive Effort. Neuron. 89, 695–710 (2016).

3. J. Berke, What does dopamine mean? Nat. Neurosci. 21, 787–793 (2018).

4. R. S. Sutton, A. G. Barto, Reinforcement Learning: An Introduction (1998; http://ieeexplore.ieee.org/document/712192/), vol. 9.

5. N. X. Tritsch, B. L. Sabatini, Dopaminergic Modulation of Synaptic Transmission in Cortex and Striatum. Neuron. 76, 33–50 (2012).

6. K. He, M. Huertas, S. Z. Hong, X. Tie, J. W. Hell, H. Shouval, A. Kirkwood, Distinct Eligibility Traces for LTP and LTD in Cortical Synapses. Neuron. 88, 528–538 (2015).

7. T. Shindou, M. Shindou, S. Watanabe, J. Wickens, A silent eligibility trace enables dopamine-dependent synaptic plasticity for reinforcement learning in the mouse striatum. Eur. J. Neurosci. 49, 726–736 (2019).

8. E. M. Izhikevich, Solving the distal reward problem through linkage of STDP and dopamine signaling. Cereb. Cortex. 17, 2443–2452 (2007).

9. Z. Brzosko, W. Schultz, O. Paulsen, Retroactive modulation of spike timing-dependent plasticity by dopamine. eLife. 4, e09685 (2015).

10. E. E. Steinberg, R. Keiflin, J. R. Boivin, I. B. Witten, K. Deisseroth, P. H. Janak, A causal link between prediction errors, dopamine neurons and learning. Nat. Neurosci. 16, 966–973 (2013).

11. A. A. Hamid, J. R. Pettibone, O. S. Mabrouk, V. L. Hetrick, R. Schmidt, C. M. Vander Weele, R. T. Kennedy, B. J. Aragona, J. D. Berke, Mesolimbic dopamine signals the value of work. Nat. Neurosci. 19, 117–126 (2016).

12. L. T. Coddington, J. T. Dudman, Learning from Action: Reconsidering Movement Signaling in Midbrain Dopamine Neuron Activity. Neuron. 104, 63–77 (2019).

13. A. Klaus, J. Alves da Silva, R. M. Costa, What, If, and When to Move: Basal Ganglia Circuits and Self-Paced Action Initiation. Annu. Rev. Neurosci. 42, 459–483 (2019).

14. M. W. Howe, D. A. Dombeck, Rapid signalling in distinct dopaminergic axons during locomotion and reward. Nature. 535, 505–510 (2016).

15. E. C. J. Syed, L. L. Grima, P. J. Magill, R. Bogacz, P. Brown, M. E. Walton, Action initiation shapes mesolimbic dopamine encoding of future rewards. Nat. Neurosci. 19, 34–36 (2016).

16. L. T. Coddington, J. T. Dudman, The timing of action determines reward prediction signals in identified midbrain dopamine neurons. Nat. Neurosci. 21, 1563–1573 (2018).

17. J. A. da Silva, F. Tecuapetla, V. Paixão, R. M. Costa, Dopamine neuron activity before action initiation gates and invigorates future movements. Nature. 554, 244–248 (2018).

18. J. W. Barter, S. Li, D. Lu, R. A. Bartholomew, M. A. Rossi, C. T. Shoemaker, D. Salas-Meza, E. Gaidis, H. H. Yin, Beyond reward prediction errors: the role of dopamine in movement kinematics. Front. Integr. Neurosci. 9, 39 (2015).

19. J. D. Salamone, M. Correa, The Mysterious Motivational Functions of Mesolimbic Dopamine. Neuron. 76, 470–485 (2012).

20. B. Engelhard, J. Finkelstein, J. Cox, W. Fleming, H. J. Jang, S. Ornelas, S. A. Koay, S. Y. Thiberge, N. Daw, D. W. Tank, I. B. Witten, Specialized coding of sensory, motor, and cognitive variables in VTA dopamine neurons. Nature. 570, 509–513 (2019).

21. S. M. Nicola, The Flexible Approach Hypothesis: Unification of Effort and Cue-Responding Hypotheses for the Role of Nucleus Accumbens Dopamine in the Activation of Reward-Seeking Behavior. J. Neurosci. 30, 16585–16600 (2010).

22. M. E. Walton, S. Bouret, What Is the Relationship between Dopamine and Effort? Trends Neurosci. 42, 79–91 (2019).

23. D. O. Hebb, The organization of behavior; a neuropsycholocigal theory. (John Wiley & Sons, Inc., New York, 1949).

24. J. J. Hopfield, Neural networks and physical systems with emergent collective computational abilities. Proc. Natl. Acad. Sci. 79, 2554–2558 (1982).

25. N. Brunel, X. J. Wang, Effects of neuromodulation in a cortical network model of object working memory dominated by recurrent inhibition. J. Comput. Neurosci. 11, 63–85 (2001).

26. X.-J. Wang, Probabilistic Decision Making by Slow Reverberation in Cortical Circuits. Neuron. 36, 955–968 (2002).

27. D. Durstewitz, G. Deco, Computational significance of transient dynamics in cortical networks. Eur. J. Neurosci. 27, 217–227 (2008).

28. D. Durstewitz, J. K. Seamans, The Dual-State Theory of Prefrontal Cortex Dopamine Function with Relevance to Catechol-O-Methyltransferase Genotypes and Schizophrenia. Biol. Psychiatry. 64, 739–749 (2008).

29. S. B. Flagel, H. Akil, T. E. Robinson, Individual differences in the attribution of incentive salience to reward-related cues: Implications for addiction. Neuropharmacology. 56, 139–148 (2009).

30. J. Naudé, S. Tolu, M. Dongelmans, N. Torquet, S. Valverde, G. Rodriguez, S. Pons, U. Maskos, A. Mourot, F. Marti, P. Faure, Nicotinic receptors in the ventral tegmental area promote uncertainty-seeking. Nat. Neurosci. 19, 471–478 (2016).

31. M. Dongelmans, R. Durand-de Cuttoli, C. Nguyen, M. Come, E. K. Duranté, D. Lemoine, R. Brito, T. Ahmed Yahia, S. Mondoloni, S. Didienne, E. Bousseyrol, B. Hannesse, L. M. Reynolds, N. Torquet, D. Dalkara, F. Marti, A. Mourot, J. Naudé, P. Faure, Chronic nicotine increases midbrain dopamine neuron activity and biases individual strategies towards reduced exploration in mice. Nat. Commun. 12, 6945 (2021).

32. H.-C. Tsai, F. Zhang, A. Adamantidis, G. D. Stuber, A. Bonci, L. de Lecea, K. Deisseroth, Phasic Firing in Dopaminergic Neurons Is Sufficient for Behavioral Conditioning. Science. 324, 1080–1084 (2009).

33. J. K. Seamans, C. R. Yang, The principal features and mechanisms of dopamine modulation in the prefrontal cortex. Prog. Neurobiol. 74, 1–58 (2004).

34. P. Cisek, Cortical mechanisms of action selection: the affordance competition hypothesis. Philos. Trans. R. Soc. B Biol. Sci. 362, 1585–1599 (2007).

35. D. R. Euston, A. J. Gruber, B. L. McNaughton, The Role of Medial Prefrontal Cortex in Memory and Decision Making. Neuron. 76, 1057–1070 (2012).

36. V. Hok, E. Save, P. P. Lenck-Santini, B. Poucet, Coding for spatial goals in the prelimbic/infralimbic area of the rat frontal cortex. Proc. Natl. Acad. Sci. 102, 4602–4607 (2005).

37. M. Rigotti, O. Barak, M. R. Warden, X.-J. Wang, N. D. Daw, E. K. Miller, S. Fusi, The importance of mixed selectivity in complex cognitive tasks. Nature. 497, 585–590 (2013).

38. M. R. Penner, S. J. Y. Mizumori, Neural systems analysis of decision making during goal-directed navigation. Prog. Neurobiol. 96, 96–135 (2012).

39. L. T. Hunt, B. Y. Hayden, A distributed, hierarchical and recurrent framework for rewardbased choice. Nat. Rev. Neurosci. 18, 172–182 (2017).

40. C. R. Gallistel, P. Shizgal, J. S. Yeomans, A portrait of the substrate for self-stimulation. Psychol. Rev. 88, 228–273 (1981).

41. Y. Niv, N. D. Daw, D. Joel, P. Dayan, Tonic dopamine: opportunity costs and the control of response vigor. Psychopharmacology (Berl.). 191, 507–520 (2007).

42. E. C. Tolman, Cognitive maps in rats and men. Psychol. Rev. 55, 189–208 (1948).

43. B. W. Balleine, The Meaning of Behavior: Discriminating Reflex and Volition in the Brain. Neuron. 104, 47–62 (2019).

44. M. Z. Wang, B. Y. Hayden, Latent learning, cognitive maps, and curiosity. Curr. Opin. Behav. Sci. 38, 1–7 (2021).

## References for Methods

1. X.-J. Wang, Probabilistic Decision Making by Slow Reverberation in Cortical Circuits. Neuron. 36, 955–968 (2002).

2. P. Cisek, Cortical mechanisms of action selection: the affordance competition hypothesis. Philos. Trans. R. Soc. B Biol. Sci. 362, 1585–1599 (2007).

3. D. R. Euston, A. J. Gruber, B. L. McNaughton, The Role of Medial Prefrontal Cortex in Memory and Decision Making. Neuron. 76, 1057–1070 (2012).

4. M. R. Penner, S. J. Y. Mizumori, Neural systems analysis of decision making during goal-directed navigation. Prog. Neurobiol. 96, 96–135 (2012).

5. L. T. Hunt, B. Y. Hayden, A distributed, hierarchical and recurrent framework for rewardbased choice. Nat. Rev. Neurosci. 18, 172–182 (2017).

6. S. M. Nicola, The Flexible Approach Hypothesis: Unification of Effort and Cue-Responding Hypotheses for the Role of Nucleus Accumbens Dopamine in the Activation of Reward-Seeking Behavior. J. Neurosci. 30, 16585–16600 (2010).

7. J. D. Salamone, M. Correa, The Mysterious Motivational Functions of Mesolimbic Dopamine. Neuron. 76, 470–485 (2012).

8. A. Arleo, W. Gerstner, Spatial cognition and neuro-mimetic navigation: a model of hippocampal place cell activity. Biol. Cybern. 83, 287–299 (2000).

9. B. Girard, N. Tabareau, Q. C. Pham, A. Berthoz, J.-J. Slotine, Where neuroscience and dynamic system theory meet autonomous robotics: A contracting basal ganglia model for action selection. Neural Netw. 21, 628–641 (2008).

10. A. Klaus, J. Alves da Silva, R. M. Costa, What, If, and When to Move: Basal Ganglia Circuits and Self-Paced Action Initiation. Annu. Rev. Neurosci. 42, 459–483 (2019).

11. K. Gurney, T. J. Prescott, P. Redgrave, A computational model of action selection in the basal ganglia. I. A new functional anatomy. Biol. Cybern. 84, 401–410 (2001).

12. M. Humphries, M. Khamassi, K. Gurney, Dopaminergic control of the explorationexploitation trade-off via the basal ganglia. Front. Neurosci. 6 (2012) (available at https://www.frontiersin.org/articles/10.3389/fnins.2012.00009).

13. N. X. Tritsch, B. L. Sabatini, Dopaminergic Modulation of Synaptic Transmission in Cortex and Striatum. Neuron. 76, 33–50 (2012).

14. K. He, M. Huertas, S. Z. Hong, X. Tie, J. W. Hell, H. Shouval, A. Kirkwood, Distinct Eligibility Traces for LTP and LTD in Cortical Synapses. Neuron. 88, 528–538 (2015).

15. Q. Cai, M. Zeng, X. Wu, H. Wu, Y. Zhan, R. Tian, M. Zhang, CaMKIIa-driven, phosphatase-checked postsynaptic plasticity via phase separation. Cell Res. 31, 37–51 (2021).

16. J. Zhang, S.-Y. Ko, Y. Liao, Y. Kwon, S. J. Jeon, A. Sohn, J. H. Cheong, D. H. Kim, J. H. Ryu, Activation of the dopamine D1 receptor can extend long-term spatial memory persistence via PKA signaling in mice. Neurobiol. Learn. Mem. 155, 568–577 (2018).

17. K. Okuda, K. Højgaard, L. Privitera, G. Bayraktar, T. Takeuchi, Initial memory consolidation and the synaptic tagging and capture hypothesis. Eur. J. Neurosci. (2020), doi:10.1111/ejn.14902.

18. J. C. Magee, C. Grienberger, Synaptic Plasticity Forms and Functions. Annu. Rev. Neurosci. 43, 95–117 (2020).

19. J. K. Seamans, C. R. Yang, The principal features and mechanisms of dopamine modulation in the prefrontal cortex. Prog. Neurobiol. 74, 1–58 (2004).

20. R. S. Sutton, A. G. Barto, Reinforcement Learning: An Introduction (1998; http://ieeexplore.ieee.org/document/712192/), vol. 9.

21. W. Schultz, Behavioral dopamine signals. Trends Neurosci. 30, 203–210 (2007).

22. K. C. Berridge, T. E. Robinson, What is the role of dopamine in reward: hedonic impact, reward learning, or incentive salience? Brain Res. Rev. 28, 309–369 (1998).

23. E. E. Steinberg, R. Keiflin, J. R. Boivin, I. B. Witten, K. Deisseroth, P. H. Janak, A causal link between prediction errors, dopamine neurons and learning. Nat. Neurosci. 16, 966–973 (2013).

24. J. Berke, What does dopamine mean? Nat. Neurosci. 21, 787–793 (2018).

25. S. M. McClure, N. D. Daw, P. Read Montague, A computational substrate for incentive salience. Trends Neurosci. 26, 423–428 (2003).

26. S. Soares, B. V. Atallah, J. J. Paton, Midbrain dopamine neurons control judgment of time. Science. 354, 1273–1277 (2016).

27. D. Durstewitz, J. K. Seamans, The Dual-State Theory of Prefrontal Cortex Dopamine Function with Relevance to Catechol-O-Methyltransferase Genotypes and Schizophrenia. Biol. Psychiatry. 64, 739–749 (2008).

28. A. Westbrook, T. S. Braver, Dopamine Does Double Duty in Motivating Cognitive Effort. Neuron. 89, 695–710 (2016).

29. J. A. da Silva, F. Tecuapetla, V. Paixão, R. M. Costa, Dopamine neuron activity before action initiation gates and invigorates future movements. Nature. 554, 244–248 (2018).

30. M. W. Howe, D. A. Dombeck, Rapid signalling in distinct dopaminergic axons during locomotion and reward. Nature. 535, 505–510 (2016).

31. A. A. Hamid, J. R. Pettibone, O. S. Mabrouk, V. L. Hetrick, R. Schmidt, C. M. Vander Weele, R. T. Kennedy, B. J. Aragona, J. D. Berke, Mesolimbic dopamine signals the value of work. Nat. Neurosci. 19, 117–126 (2016).

32. L. T. Coddington, J. T. Dudman, The timing of action determines reward prediction signals in identified midbrain dopamine neurons. Nat. Neurosci. 21, 1563–1573 (2018).

33. L. T. Coddington, J. T. Dudman, Learning from Action: Reconsidering Movement Signaling in Midbrain Dopamine Neuron Activity. Neuron. 104, 63–77 (2019).

34. E. C. J. Syed, L. L. Grima, P. J. Magill, R. Bogacz, P. Brown, M. E. Walton, Action initiation shapes mesolimbic dopamine encoding of future rewards. Nat. Neurosci. 19, 34–36 (2016).

35. X.-J. Wang, Decision Making in Recurrent Neuronal Circuits. Neuron. 60, 215–234 (2008).

36. I. Tsuda, Toward an interpretation of dynamic neural activity in terms of chaotic dynamical systems. Behav. Brain Sci. 24, 793–810 (2001).

37. G. Deco, V. K. Jirsa, P. A. Robinson, M. Breakspear, K. Friston, The Dynamic Brain: From Spiking Neurons to Neural Masses and Cortical Fields. PLOS Comput. Biol. 4, e1000092 (2008).

38. B. W. Balleine, The Meaning of Behavior: Discriminating Reflex and Volition in the Brain. Neuron. 104, 47–62 (2019).

39. M. K. Muezzinoglu, I. Tristan, R. Huerta, V. S. Afraimovich, M. I. Rabinovich, Transients versus attractors in complex networks. Int. J. Bifurc. Chaos. 20, 1653–1675 (2010).

40. C. Gros, Neural networks with transient state dynamics. New J. Phys. 9, 109–109 (2007).

41. E. C. Tolman, Cognitive maps in rats and men. Psychol. Rev. 55, 189–208 (1948).

42. M. Z. Wang, B. Y. Hayden, Latent learning, cognitive maps, and curiosity. Curr. Opin. Behav. Sci. 38, 1–7 (2021).

43. M. Turiault, S. Parnaudeau, A. Milet, R. Parlato, J.-D. Rouzeau, M. Lazar, F. Tronche, Analysis of dopamine transporter gene expression pattern - generation of DAT-iCre transgenic mice. FEBS J. 274, 3568–3577 (2007).

44. N. Benamer, F. Marti, R. Lujan, R. Hepp, T. G. Aubier, A. a. M. Dupin, G. Frébourg, S. Pons, U. Maskos, P. Faure, Y. A. Hay, B. Lambolez, L. Tricoire, GluD1, linked to schizophrenia, controls the burst firing of dopamine neurons. Mol. Psychiatry. 23, 691–700 (2018).

45. H. Khabou, M. Garita-Hernandez, A. Chaffiol, S. Reichman, C. Jaillard, E. Brazhnikova, S. Bertin, V. Forster, M. Desrosiers, C. Winckler, O. Goureau, S. Picaud, J. Duebel, J.-A. Sahel, D. Dalkara, Noninvasive gene delivery to foveal cones for vision restoration. JCI Insight. 3, 96029 (2018).

46. V. W. Choi, A. Asokan, R. A. Haberman, R. J. Samulski, Curr. Protoc. Mol. Biol., in press, doi:10.1002/0471142727.mb1625s78.

47. C. Aurnhammer, M. Haase, N. Muether, M. Hausl, C. Rauschhuber, I. Huber, H. Nitschko, U. Busch, A. Sing, A. Ehrhardt, A. Baiker, Universal Real-Time PCR for the Detection and Quantification of Adeno-Associated Virus Serotype 2-Derived Inverted Terminal Repeat Sequences. Hum. Gene Ther. Methods. 23, 18–28 (2012).

48. M. X. B. Sarazin, J. Victor, D. Medernach, J. Naudé, B. Delord, Online Learning and Memory of Neural Trajectory Replays for Prefrontal Persistent and Dynamic Representations in the Irregular Asynchronous State. Front. Neural Circuits. 0 (2021), doi:10.3389/fncir.2021.648538.

49. E. M. Izhikevich, Solving the distal reward problem through linkage of STDP and dopamine signaling. Cereb. Cortex. 17, 2443–2452 (2007).

50. M. Graupner, N. Brunel, Calcium-based plasticity model explains sensitivity of synaptic changes to spike pattern, rate, and dendritic location. Proc. Natl. Acad. Sci., 201109359 (2012).

## References for supplementary methods

1. E. Todorov, M. I. Jordan, Optimal feedback control as a theory of motor coordination. Nat. Neurosci. 5, 1226–1235 (2002).

2. D. J. Best, N. I. Fisher, Efficient Simulation of the von Mises Distribution. J. R. Stat. Soc. Ser. C Appl. Stat. 28, 152–157 (1979).

3. M. X. B. Sarazin, J. Victor, D. Medernach, J. Naudé, B. Delord, Online Learning and Memory of Neural Trajectory Replays for Prefrontal Persistent and Dynamic Representations in the Irregular Asynchronous State. Front. Neural Circuits. 0 (2021), doi: 10.3389/fncir.2021.648538.

4. H. Dale, Pharmacology and Nerve-endings (Walter Ernest Dixon Memorial Lecture): (Section of Therapeutics and Pharmacology). Proc. R. Soc. Med. 28, 319–332 (1935).

5. C. Beaulieu, Z. Kisvarday, P. Somogyi, M. Cynader, A. Cowey, Quantitative distribution of gaba-immunopositive and-immunonegative neurons and synapses in the monkey striate cortex (area 17). Cereb. Cortex. 2, 295–309 (1992).

6. A. M. Thomson, Synaptic Connections and Small Circuits Involving Excitatory and Inhibitory Neurons in Layers 2-5 of Adult Rat and Cat Neocortex: Triple Intracellular Recordings and Biocytin Labelling In Vitro. Cereb. Cortex. 12, 936–953 (2002).

7. N. Brunel, X. J. Wang, Effects of neuromodulation in a cortical network model of object working memory dominated by recurrent inhibition. J. Comput. Neurosci. 11, 63–85 (2001).

8. C. E. Jahr, C. F. Stevens, Voltage dependence of NMDA-activated macroscopic conductances predicted by single-channel kinetics. J. Neurosci. 10, 3178–3182 (1990).

9. G. Chen, P. Greengard, Z. Yan, Potentiation of NMDA receptor currents by dopamine D1 receptors in prefrontal cortex. Proc. Natl. Acad. Sci. 101, 2596–2600 (2004).

10. J. K. Seamans, C. R. Yang, The principal features and mechanisms of dopamine modulation in the prefrontal cortex. Prog. Neurobiol. 74, 1–58 (2004).

11. N. X. Tritsch, B. L. Sabatini, Dopaminergic Modulation of Synaptic Transmission in Cortex and Striatum. Neuron. 76, 33–50 (2012).

12. H. Wang, G. G. Stradtman, X.-J. Wang, W.-J. Gao, A specialized NMDA receptor function in layer 5 recurrent microcircuitry of the adult rat prefrontal cortex. Proc. Natl. Acad. Sci. (2008), doi:10.1073/pnas.0804318105.

13. A. Destexhe, Z. F. Mainen, T. J. Sejnowski, Kinetic models of synaptic transmission. Methods Neuronal Model. 2, 1–25 (1998).

14. M. Xue, B. V. Atallah, M. Scanziani, Equalizing excitation–inhibition ratios across visual cortical neurons. Nature. 511, 596–600 (2014).

15. R. S. Sutton, A. G. Barto, Reinforcement Learning: An Introduction (1998; http://ieeexplore.ieee.org/document/712192/), vol. 9.

16. E. M. Izhikevich, Solving the distal reward problem through linkage of STDP and dopamine signaling. Cereb. Cortex. 17, 2443–2452 (2007).

17. G. Q. Bi, M. M. Poo, Synaptic modifications in cultured hippocampal neurons: dependence on spike timing, synaptic strength, and postsynaptic cell type. J. Neurosci. Off. J. Soc. Neurosci. 18, 10464–10472 (1998).

18. K. He, M. Huertas, S. Z. Hong, X. Tie, J. W. Hell, H. Shouval, A. Kirkwood, Distinct Eligibility Traces for LTP and LTD in Cortical Synapses. Neuron. 88, 528–538 (2015).

19. M. Graupner, N. Brunel, Calcium-based plasticity model explains sensitivity of synaptic changes to spike pattern, rate, and dendritic location. Proc. Natl. Acad. Sci., 201109359 (2012).

20. T. Shindou, M. Shindou, S. Watanabe, J. Wickens, A silent eligibility trace enables dopamine-dependent synaptic plasticity for reinforcement learning in the mouse striatum. Eur. J. Neurosci. 49, 726–736 (2019).

21. J. C. Magee, C. Grienberger, Synaptic Plasticity Forms and Functions. Annu. Rev. Neurosci. 43, 95–117 (2020).

22. N. Frémaux, W. Gerstner, Neuromodulated Spike-Timing-Dependent Plasticity, and Theory of Three-Factor Learning Rules. Front. Neural Circuits. 9 (2016), doi:10.3389/fncir.2015.00085.

23. J. Zhang, S.-Y. Ko, Y. Liao, Y. Kwon, S. J. Jeon, A. Sohn, J. H. Cheong, D. H. Kim, J. H. Ryu, Activation of the dopamine D1 receptor can extend long-term spatial memory persistence via PKA signaling in mice. Neurobiol. Learn. Mem. 155, 568–577 (2018).

24. K. Okuda, K. Højgaard, L. Privitera, G. Bayraktar, T. Takeuchi, Initial memory consolidation and the synaptic tagging and capture hypothesis. Eur. J. Neurosci. (2020), doi:10.1111/ejn.14902.

25. Z. Brzosko, W. Schultz, O. Paulsen, Retroactive modulation of spike timing-dependent plasticity by dopamine. eLife. 4, e09685 (2015).

26. D. Muir, vmrand(fMu, fKappa, varargin), version 1.3.0.0. MATLAB Cent. File Exch. (2017), (available at https://www.mathworks.com/matlabcentral/fileexchange/37241-vmrand-fmu-fkappa-varargin).

27. S. Strogatz, M. Friedman, A. J. Mallinckrodt, S. McKay, Nonlinear Dynamics and Chaos: With Applications to Physics, Biology, Chemistry, and Engineering. Comput. Phys. 8, 532–532 (1994).

